# Quantifying the impact of a periodic presence of antimicrobial on resistance evolution in a homogeneous microbial population of fixed size

**DOI:** 10.1101/279091

**Authors:** Loïc Marrec, Anne-Florence Bitbol

## Abstract

The evolution of antimicrobial resistance often occurs in a variable environment, as antimicrobial is given periodically to a patient or added and removed from a medium. This environmental variability has a huge impact on the microorganisms’ fitness landscape, and thus on the evolution of resistance. Indeed, mutations conferring resistance often carry a fitness cost in the absence of antimicrobial, which may be compensated by subsequent mutations. As antimicrobial is added or removed, the relevant fitness landscape thus switches from a fitness valley to an ascending landscape or vice-versa.

Here, we investigate the effect of these time-varying patterns of selection within a stochastic model. We focus on a homogeneous microbial population of fixed size subjected to a periodic alternation of phases of absence and presence of an antimicrobial that stops growth. Combining analytical approaches and stochastic simulations, we quantify how the time necessary for fit resistant bacteria to take over the microbial population depends on the period of the alternations. We demonstrate that fast alternations strongly accelerate the evolution of resistance, and that a plateau is reached once the period gets sufficiently small. Besides, the acceleration of resistance evolution is stronger for larger populations. For asymmetric alternations, featuring a different duration of the phases with and without antimicrobial, we shed light on the existence of a broad minimum of the time taken by the population to fully evolve resistance. At this minimum, if the alternations are sufficiently fast, the very first resistant mutant that appears ultimately leads to full resistance evolution within the population. This dramatic acceleration of the evolution of antimicrobial resistance likely occurs in realistic situations, and can have an important impact both in clinical and experimental situations.

## Introduction

The discovery of antibiotics and antivirals has constituted one of the greatest medical advances of the twentieth century, allowing many major infectious diseases to be treated. However, with the increasing use of antimicrobials, pathogenic microorganisms tend to become resistant to these drugs. Antimicrobial resistance has become a major and urgent problem of public health worldwide [1, 2].

Mutations that confer antimicrobial resistance are often associated with a fitness cost, i.e. a slower reproduction [3, 4]. Indeed, the acquisition of resistance generally involves either a modification of the molecular target of the antimicrobial, which often alters its biological function, or the production of specific proteins, which entails a metabolic cost [5]. However, resistant microorganisms frequently acquire subsequent mutations that compensate for the initial cost of resistance. These microorganisms are called “resistant-compensated” [6, 7, 8, 9]. The acquisition of resistance is therefore often irreversible, even if the antimicrobial is removed from the environment [10, 4].

In the absence of antimicrobial, the adaptive landscape of the microorganism, which represents its fitness (i.e. its reproduction rate) as a function of its genotype, involves a valley, since the first resistance mutation decreases fitness, while compensatory mutations increase it. However, this fitness valley, which exists in the absence of antimicrobial, disappears above a certain concentration of antimicrobial, as the growth of the antimicrobial-sensitive microorganism is impaired. Thus, the adaptive landscape of the microorganism depends drastically on whether the antimicrobial is present or absent. Recent experiments show that this type of interaction between genotype and environment is common and important [11, 12]. Taking into account these effects constitutes a fundamental problem, which has been little studied so far, particularly because experimental works have traditionally focused on comparing different mutants in a unique environment [13]. However, recent theoretical analyses show that variable adaptive landscapes can have a dramatic evolutionary impact [14, 15, 16, 17, 18, 19].

How do the time scales of the evolution and of the variation of the adaptive landscape couple together? What is the impact of the time variability of the adaptive landscape on the evolution of antimicrobial resistance? In order to answer these questions, we construct a minimal model retaining the fundamental aspects of antimicrobial resistance evolution. Focusing on the case of a homogeneous microbial population of fixed size, we perform a complete stochastic study of *de novo* resistance acquisition in the presence of alternations of phases of absence and presence of an antimicrobial that stops growth. These alternations can represent, for example, a treatment where the concentration within the patient falls under the Minimum Inhibitory Concentration (MIC) between drug intakes [20]. Combining analytical and numerical approaches, we show that these alternations substantially accelerate the evolution of resistance, especially for larger populations. We fully quantify this effect and shed light on the different regimes at play. We also compare the alternation-driven acquisition of resistance to the spontaneous evolution of resistance by fitness valley crossing, and extend previous results on valley crossing [21]. We then generalize our study to the case of asymmetric alternations, featuring a different duration of the phases with and without antimicrobial. We demonstrate the existence of a broad minimum of the time taken by the population to fully evolve resistance, occurring when both phases have durations of the same order. This realistic situation dramatically accelerates the evolution of resistance. Finally, we discuss the implications of our findings, in particular regarding antimicrobial dosage.

## Model

The action of an antimicrobial drug can be quantified by its MIC, the minimum concentration that stops the growth of a microbial population [4]. We focus on biostatic antimicrobials, which stop microbial growth (vs. biocidal antimicrobials, which kill microorganisms). We model the action of the antimicrobial in a binary way: below the MIC (“absence of antimicrobial”), growth is not affected, while above it (“presence of antimicrobial”), sensitive microorganisms cannot grow at all. The usual stiffness of pharmacodynamic curves around the MIC [20] justifies our simple binary approximation, and we also present an analysis of the robustness of this hypothesis (see S1 Appendix). Within this binary approximation, there are two adaptive landscapes, which can be described minimally by a single parameter *δ*, representing the fitness cost of resistance (Fig. 1A). We focus on asexual microorganisms, and fitness simply denotes the division rate of these organisms. The fitness of sensitive microorganisms in the absence of antimicrobials is taken as reference, and is used as the time unit throughout. In this framework, we investigate the impact of a periodic presence of antimicrobial, assuming that the process starts without antimicrobial (Fig. 1B-C).

**Figure 1:**
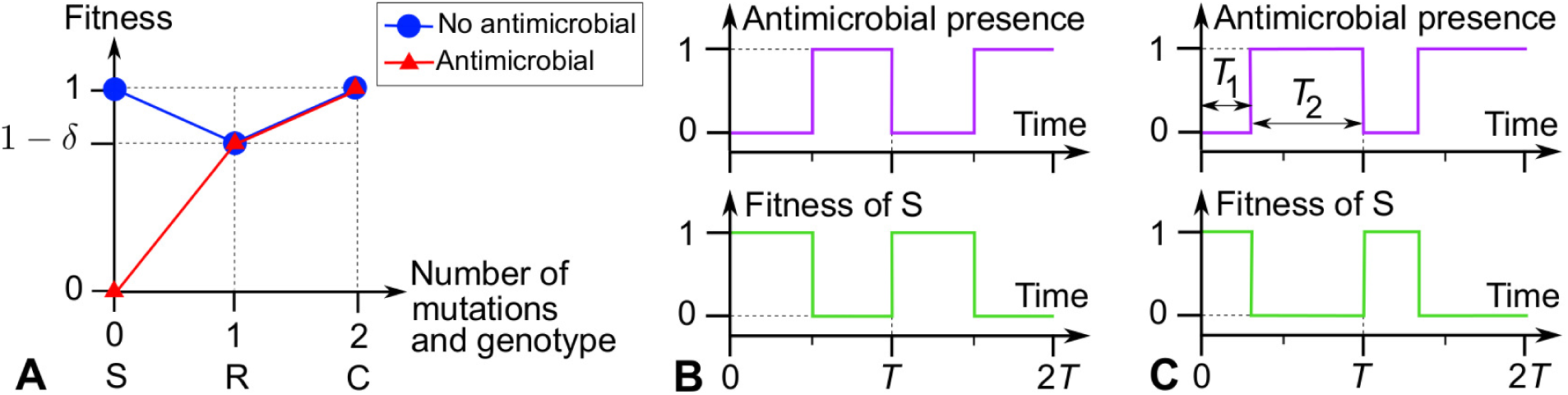
Model. (A) Adaptive landscapes in the presence and in the absence of antimicrobial. Genotypes are indicated by the number of mutations from the sensitive microorganism, and by initials: S: sensitive; R: resistant; C: resistant-compensated. (B) and (C) Periodic presence of antimicrobial, and impact on the fitness of S (sensitive) microorganisms: (B) Symmetric alternations; (C) Asymmetric alternations.

We denote by *µ*_1_ and *µ*_2_ the mutation rates (or mutation probabilities upon each division) for the mutation from S to R and for the one from R to C, respectively. In several actual situations, the effective mutation rate towards compensation tends to be higher than the one towards the return to sensitivity, since multiple mutations can compensate for the initial cost of resistance [7, 8, 22]. Therefore, we do not take into account back-mutations. Still because of the abundance of possible compensatory mutations, generically *µ*_1_ *≪µ*_2_ [7, 23]. We present general analytical results as a function of *µ*_1_ and *µ*_2_, and analyze in more detail the limit *µ*_1_ *≪µ*_2_, especially in simulations. All notations introduced are summed up in Table S1.

We focus on a homogeneous microbial population of fixed size *N*, which can thus be described in the framework of the Moran process [24, 25] (see S1 Appendix and Fig. S1). This simplifying hypothesis is appropriate to describe an experiment in a chemostat [26], and should constitute a reasonable approximation in the intermediate stages of an infection, sufficiently after its onset and before its eradication. Within the Moran process, fitnesses are relative. If a population only features sensitive individuals (with zero fitness) in the presence of antimicrobial, we consider that no division occurs, and the population remains static. We always express time in number of generations, which corresponds (unless no cell can divide) to the number of Moran steps divided by the population size *N*, since each Moran step corresponds to the division of one individual and the death of one individual

Throughout, we start from a microbial population where all individuals are S (sensitive), and we focus on the time 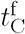 it takes for the C (resistant-compensated) type to fix in the population, i.e. to take over the population. Then, all individuals are resistant-compensated, and the population has fully evolved resistance *de novo*. The main originality of our model arises from the variability of the adaptive landscape. We compare the different environmental and evolutionary timescales, and make analytical predictions for the average time needed for the population to fully evolve resistance in each regime where timescales are separated. Numerical simulations of the Moran process allow us to test our analytical predictions, and to go beyond separated timescales.

## Results

### A periodic presence of antimicrobial can drive resistance evolution

In this section, we demonstrate how alternations of absence and presence of antimicrobial can drive the *de novo* evolution of resistance. We first present analytical predictions for the time needed for resistant microorganisms to start growing, and then for the total time needed for the population to fully evolve resistance. We then compare these predictions to numerical simulation results.

We first focus on the rare mutation regime *Nµ*_1_ *≪*1, where at most one mutant lineage exists in the population at each given time, and there is no clonal interference. The frequent mutation regime is briefly discussed, and more detail regarding the appropriate deterministic treatment in this regime is given in S1 Appendix. Here, we consider the case of symmetric alternations with period *T* (see Fig. 1B). The more general case of asymmetric alternations (Fig. 1C) is discussed later.

### Time needed for resistant microorganisms to start growing

Resistant (R) mutants can only appear in the absence of antimicrobial. Indeed, mutations occur upon division, and sensitive (S) bacteria cannot divide in the presence of antimicrobial (as their fitness is zero, see Fig. 1). During phases without antimicrobial, R individuals feature a lower fitness (1*-δ*) than S individuals (which then have fitness 1), see Fig. 1A. Hence, the lineage of an R mutant will very likely disappear, unless it survives until the next addition of antimicrobial. More precisely, in the absence of antimicrobial, the fixation probability *p*_SR_ of a single R mutant, in a population of size *N* where all other individuals are of type S, is *∼* 1*/N* if the mutation from S to R is effectively neutral (*Nδ ≪*1), and *∼ δe*^*-Nδ*^ if *δ ≪*1 and *Nδ ≫* 1 [25] (see S1 Appendix for derivations). The average time *τ* it takes for the R lineage to disappear is equal to the fixation time of the S type in the population, which is known for the Moran process [25] and can be recovered using the general formalism of first-passage times (see S1 Appendix). If antimicrobial is added while R mutants exist in the population, i.e. within 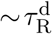 after a mutation event, then the R population will grow fast, since S individuals cannot divide with antimicrobial (see Fig. 1). Next, C individuals will be able to appear by mutation. It is thus crucial to calculate the average time 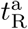 where R individuals first exist in the presence of antimicrobial, since this constitutes the first step of resistance takeover.

Three key timescales impact 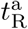. Two of them are intrinsic timescales of the evolution of the population in the absence of antimicrobial: the average time between the appearance of two independent R mutants, 1*/*(*Nµ*_1_), and the average lifetime 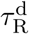 of the lineage of an R mutant before it disappears. The third one is the timescale of the environment, namely the half-period *T/*2. If mutations are sufficiently rare, 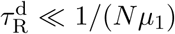. Indeed, if the mutation from S to R is neutral (i.e. *δ* = 0, see Fig. 1A), then 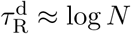 for large *N* [25] (see S1 Appendix), and 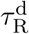 is even shorter for *δ >* 0, as deleterious R mutants are out-competed by S microorganisms. What matters is how the environment timescale *T/*2 compares to these two evolution timescales (see Fig. 2A-C).

**Figure 2:**
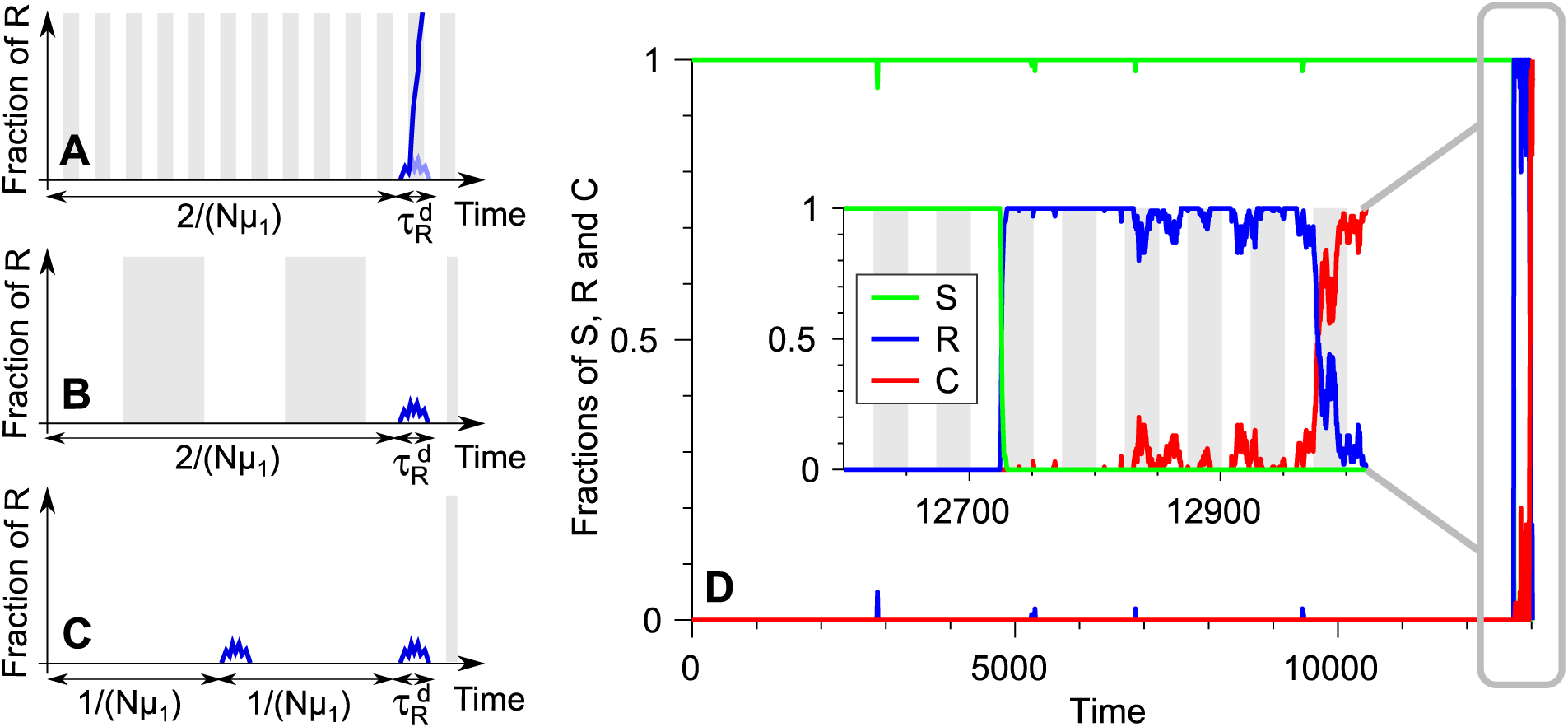
Alternation-driven evolution of antimicrobial resistance. (A-C) Sketches illustrating the three different regimes for the half-period *T/*2 of the alternations of antimicrobial absence (white) and presence (gray). The fraction of resistant (R) microorganisms in the population is plotted versus time (blue curves). R mutants can only appear without antimicrobial.

(A) *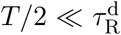*, where 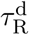 is the average extinction time of the lineage of an R mutant without antimicrobial. The first R lineage that appears lives until the next addition of antimicrobial and grows.

(B) *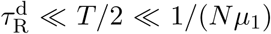*, where 1*/*(*Nµ*_1_) is the average time between the apparition of two independent R mutants without antimicrobial. (C) *T/*2 *≫* 1*/*(*Nµ*_1_). In (B) and (C), not all R lineages live until the next addition of antimicrobial, and in (C) multiple R lineages arise within a half-period. (D) Example of a simulation run. The fractions of S, R and C microorganisms are plotted versus time. Inset: end of the process, with full resistance evolution. As in (A-C), antimicrobial is present during the gray-shaded time intervals (shown only in the inset given their duration). Parameters: *µ*_1_ = 10^−5^, *µ*_2_ = 10^−3^, *δ* = 0.1, *N* = 10^2^ and *T* = 50 (belonging to regime B).

(A) If 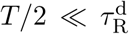 (Fig. 2A): The lineage of the very first R individual that appears will still exist upon the next addition of antimicrobial. Hence, 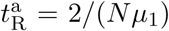. Indeed, *Nµ*_1_ represents the total mutation rate in the population, and mutation can only occur in the absence of antimicrobial, i.e. half of the time.

(B) If 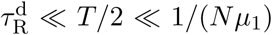 (Fig. 2B): At most one mutation yielding an R individual is expected within each half-period. The lineage of this mutant is likely to survive until the next addition of antimicrobial only if the mutant appeared within the last 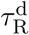 preceding it, which has a probability 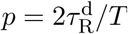. Hence, 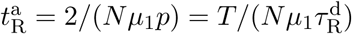.

(C) If *T/*2 *≫* 1*/*(*Nµ*_1_) (Fig. 2C): Since the half-period is much larger than the time 1*/*(*Nµ*_1_) between the appearance of two independent mutants without antimicrobial, several appearances and extinctions of R lineages are expected within one half-period. Hence, the probability that a lineage of R exists upon a given addition of antimicrobial is 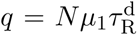. Since additions of antimicrobial occur every *T*, we have 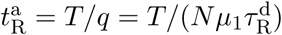, which is the same as in case (B).

In conclusion, we obtain

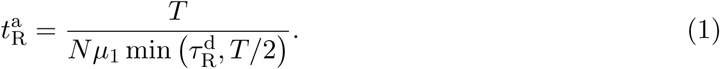

Hence, if 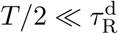, 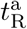 is independent from the period *T* of alternations, while if 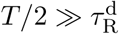, 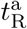 is proportional to *T*.

### Time needed for the population to fully evolve resistance

We are interested in the average time 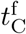 it takes for the population to fully evolve resistance, i.e.for the C (resistant-compensated) type to fix. An example of the process is shown in Fig. 2D. First, it takes on average 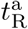 for the R mutants to first encounter antimicrobial. Then they rapidly grow, since S individuals cannot divide. If the phase with antimicrobial is long enough, R mutants take over with an average fixation timescale 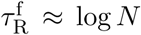 for *N ≫* 1 [25] (see S1 Appendix). If 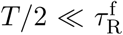, fixation cannot occur within a single half-period, and the R lineage will drift longer, but its extinction remains very unlikely. Indeed, R possesses a huge fitness advantage compared to S in the presence of antimicrobial since S cannot divide, and a smaller disadvantage without antimicrobial (1 *- δ* vs. 1, generally with *δ ≪*1 [4], see Fig. 1A). Hence, if 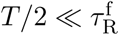 mutants will take 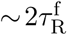 to fix. Note that for asymmetric alternations, if phases without antimicrobial are much longer than those with antimicrobial, i.e. *T*_1_ *≫ T*_2_ (see Fig. 1C), an R lineage that starts growing can go extinct (see below). But in the present case of symmetric alternations, R will fix shortly after 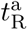, in a time of order 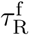.

Once the R type has fixed in the population, the appearance and eventual fixation of C mutants are independent from the presence of antimicrobial, since only S microorganisms are affected by it (see Fig. 1A). The first C mutant whose lineage will fix takes an average time 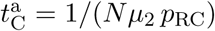 to appear once R has fixed, where *p*_RC_ is the fixation probability of a single C mutant in a population of size *N* where all other individuals are of type R. In particular, if *Nδ ≪*1 then *p*_RC_ = 1*/N*, and if *δ ≪*1 and *Nδ ≫* 1 then *p*_RC_ *≈ δ* [25] (see S1 Appendix). The final step is the fixation of this successful C mutant, which will take an average time 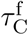, of order *N* in the effectively neutral regime *Nδ ≪*1, and shorter for larger *δ* given the selective advantage of C over R [25] (see S1 Appendix). Note that we have assumed for simplicity that the fixation of R occurs before the appearance of the first successful C mutant, which is true if 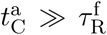, i.e. 1*/*(*Nµ*_2_ *p*_RC_) *≫* log *N*. This condition is satisfied if the second mutation is sufficiently rare. Otherwise, our calculation will slightly overestimate the actual result.

Assembling the previous results yields

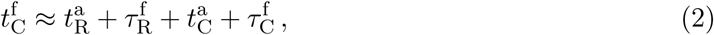

where 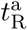 is given by Eq. 1, while, 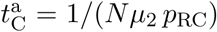 and 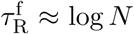 and 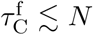 In the rare mutation regime, the contribution of the two fixation times 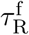 and 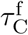 will be negligible. If in addition *µ*_1_ *≪µ*_2_, which is realistic (cf. Methods), then 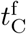 will be dominated by 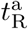 If *µ*_1_ *≈ µ*_2_, 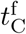 will be dominated by 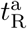 if *T >* max 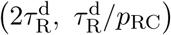, and otherwise, the contribution of 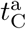 will be important.

### Comparison of analytical predictions and simulation results

Fig. 3A shows simulation results for the average total fixation time 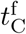 of C individuals in the population. This time is plotted as a function of the period *T* of alternations for different population sizes *N*. As predicted above (see Eq. 1), we observe two regimes delimited by 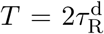 If 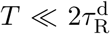, 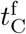 does not depend on *T*, while if 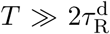, it depends linearly on *T* In Fig. 3A, we also plot our analytical prediction from Eqs. 1 and 2 in these two regimes (solid lines). The agreement with our simulated data is excellent for small and intermediate values of *T*, without any adjustable parameter.

**Figure 3:**
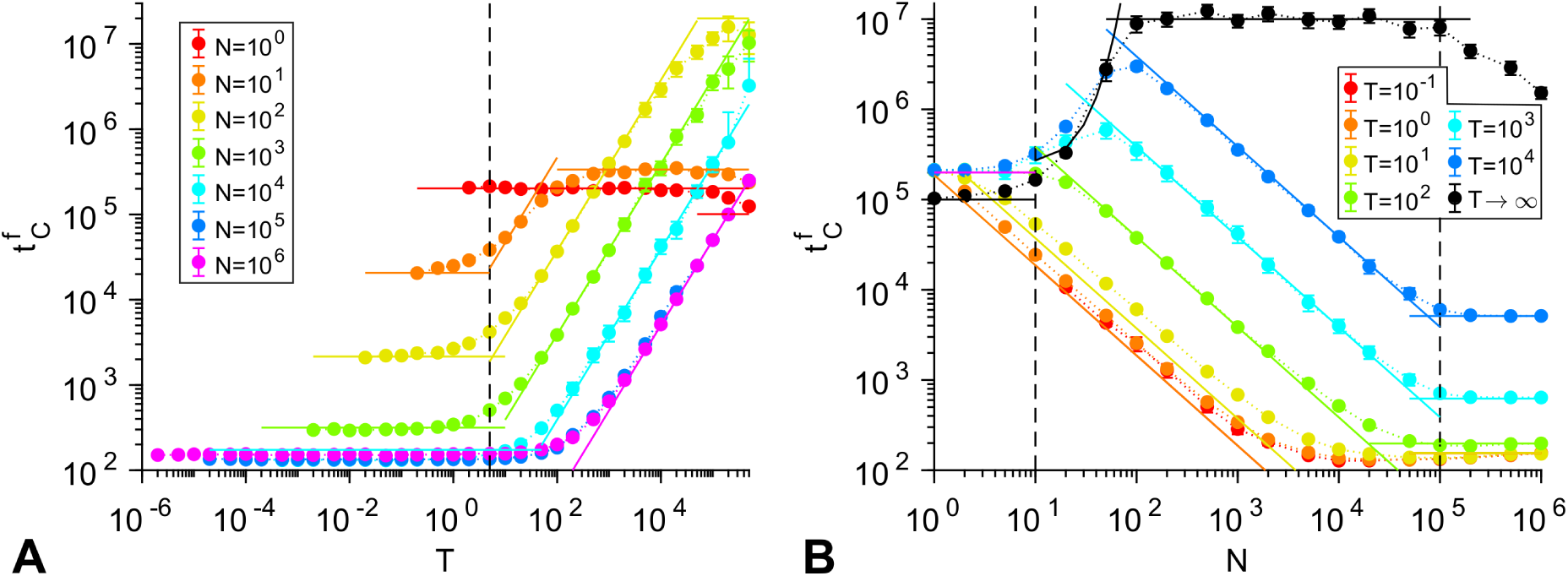
Impact of symmetric alternations. Fixation time 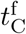 of C (resistant-compensated) individuals in a population of *N* individuals subjected to symmetric alternations of absence and presence of antimicrobial with period *T*. Data points correspond to the average of simulation results, and error bars (often smaller than markers) represent 95% confidence intervals. In both panels, solid lines correspond to our analytical predictions in each regime. Parameter values: *µ*_1_ = 10^−5^, *µ*_2_ = 10^−3^, and *δ* = 0.1. (A) 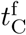 as function of *T*. Vertical dashed line: 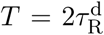. (B) 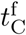 as function of *N*. Left vertical dashed line: limit of the neutral regime, *N* = 1*/δ*. Right vertical dashed line: limit of the deterministic regime, *N* = 1*/µ*_1_. Horizontal purple line:analytical prediction for valley crossing by neutral tunneling in the presence of alternations.

Importantly, Fig. 3A shows that 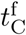 reaches a plateau for small *N* and large *T*, which cannot be predicted by our analysis of the alternation-driven evolution of resistance. This plateau corresponds to the spontaneous fitness valley crossing process [21], through which resistance can evolve without any antimicrobial. What ultimately matters is the shortest process among the alternation-driven one and the spontaneous valley-crossing one. The valley crossing time [21] can be predicted too (see below). In Fig. 3A, horizontal solid lines at large *T* correspond to these predictions.

Fig. 3B shows simulation results for 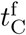 as function of *N* for different *T*. Again, solid lines represent our analytical predictions from Eqs. 1 and 2, and we obtain an excellent agreement for intermediate values of *N*, and for small ones at small *T*. In the regime of small *N* and nonsmall *T*, resistance evolution is achieved by spontaneous valley crossing, with different behaviors depending whether we are in the neutral regime *N ≪*1*/δ* or not (see below). In the limit *T → ∞* of a continuous absence of antimicrobial (black data points in Fig. 3B), only valley crossing can occur, and the black solid lines correspond to our analytical predictions for this process (see below).

Until now, we focused on the rare mutation regime. In the opposite frequent mutation regime *N ≫* 1*/µ*_1_, the dynamics of the population can be well-approximated by a deterministic model with replicator-mutator differential equations [27, 28] (see S1 Appendix). Then, clonal interference occurs, i.e. several lineages of mutants can coexist. If *T/*2 *≫* 1*/*(*Nµ*_1_), it is then almost certain that some R mutants exist in the population upon the first addition of antimicrobial, which entails 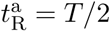. In Fig. 3A, we extended our simulations to this regime, and the horizontal purple solid line plotted at large *T* corresponds to this deterministic prediction. Similarly, in Fig. 3B, the horizontal solid lines at large *N* correspond to this deterministic prediction. In S1 Appendix, we present a derivation and a detailed study the deterministic limit of our stochastic model. We demonstrate that numerical resolution of the associated differential equations and analytical approximations match the results obtained in Fig. 3A for *N* = 10^5^ and *N* = 10^6^ over the whole range of *T*. Results are presented in Fig. S2.

Finally, note that the case *N* = 1 is special, since fixation of any mutant is immediate: 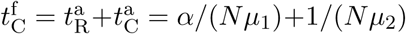, where *α* = 2 if *T/*2 *≪*1*/*(*Nµ*_1_) and *α* = 1 if *T/*2 *≫* 1*/*(*Nµ*_1_). The solid lines for *N* = 1 in Fig. 3A correspond to this prediction.

Given the usual stiffness of pharmacodynamic curves [20], we have modeled the action of the antimicrobial in a binary way, with no growth inhibition under the MIC and full growth inhibition of S microorganisms above it (see Model). An analysis of the robustness of this approximation is presented in S1 Appendix, showing that it is appropriate if the rise time Θ of the fitness satisfies 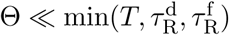 (see Fig. S3).

### Asymmetric alternations

We now turn to the more general case of asymmetric alternations of phases of absence and presence of antimicrobial, with respective durations *T*_1_ and *T*_2_, the total alternation period being *T* = *T*_1_ + *T*_2_ (see Fig. 1C).

The average time 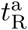 when R mutants first exist in the presence of antimicrobial, and start growing, can be obtained by a straightforward generalization of the symmetric alternation case Eq. 1. Briefly, what matters is how the duration *T*_1_ of the phase without antimicrobial, where S individuals can divide, compares to the average lifetime 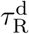 of a lineage of R mutants before extinction. We obtain

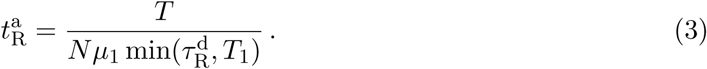

Next, we examine whether R mutants will fix during a single phase with antimicrobial, of duration *T*_2_. The fixation time of the lineage of an R mutant in the presence of antimicrobial is 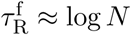 [25] for *N ≫* 1 (see S1 Appendix). If 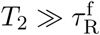, fixation will happen within *T*_2_. In the opposite case, the fixation of R is not likely to occur within a single phase with antimicrobial.Two situations exist in this case (see Fig. 4).

**Figure 4:**
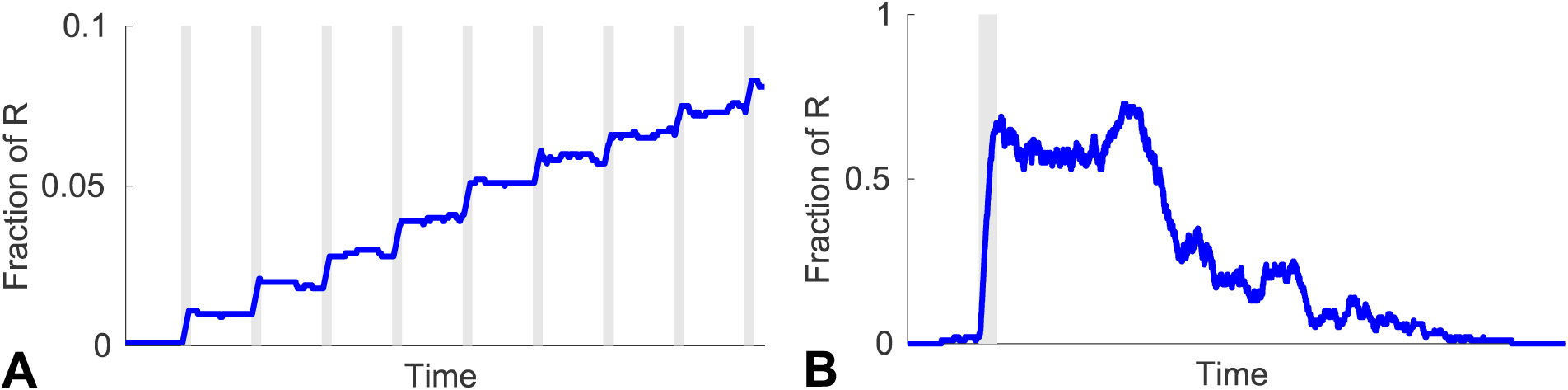
Particular regimes. The number of R individuals in the population is plotted versus time under alternations of phases without (white) and with antimicrobial (gray). Data extracted from simulation runs. (A) 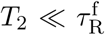 and 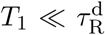 : the R lineage drifts for multiple periods. Parameters: *N* = 10^3^, *T*_1_ = 10^−1^, *T*_2_ = 10^−2^. (B) 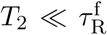 and 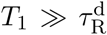 : the R lineage goes extinct. Parameters: *N* = 10^2^, *T*_1_ = 10^2^, *T*_2_ = 1. In both (A) and (B), *µ*_1_ = 10^−5^, *µ*_2_ = 10^−3^ and *δ* = 0.1.

(A) If 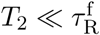 and 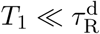 (Fig. 4A): The R lineage will drift for multiple periods, but its extinction is unlikely, as for symmetric alternations. This effect can induce a slight increase of the total time of resistance evolution, which is usually negligible.

(B) If 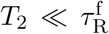 and 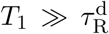 (Fig. 4B): The R lineage is likely to go extinct even after it has started growing in the presence of antimicrobial. This typically implies *T*_1_ *≫ T*_2_, since 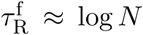 and 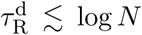 for *N ≫* 1 (see S1 Appendix). Hence, this case is specific to (very) asymmetric alternations. Spontaneous valley crossing then becomes the fastest process of resistance evolution (see below).

Once the R mutants have taken over the population, the appearance and fixation of C mutants is not affected by the alternations. Hence, Eq. 2 holds for asymmetric alternations, with 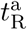 given by Eq. 3, except in the regime where 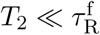 and 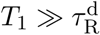 (Fig. 4B). In the rare mutation regime, if *µ*_1_ *≪µ*_2_, then 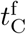 will be dominated by 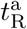, and if *u*_1_ ˜ *u*_2_, then 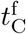 will be dominated by 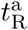 if 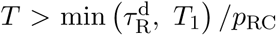, where *p*_RC_ is the fixation probability of a single C mutant in a population of R individuals.

Fig. 5A shows simulation results for 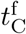 as a function of the duration *T*_1_ of the phases without antimicrobial, for different values of the duration *T*_2_ of the phases with antimicrobial. As predicted above, we observe a transition at 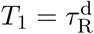, and different behaviors depending whether 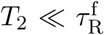 or 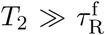. Our analytical predictions from Eqs. 2 and 3 are plotted in Fig. 5A in the various regimes (solid lines), and are in excellent agreement with the simulation data. A plateau of 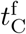 is reached at large *T*_1_, due to spontaneous valley crossing. The corresponding analytical prediction (see below) is plotted in black in Fig. 5A. Note that the large-*T*_1_ valleycrossing plateau is reached rapidly if 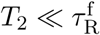, because R lineages then tend to go extinct, even once they have started growing while antimicrobial was present (see Fig. 4B).

**Figure 5:**
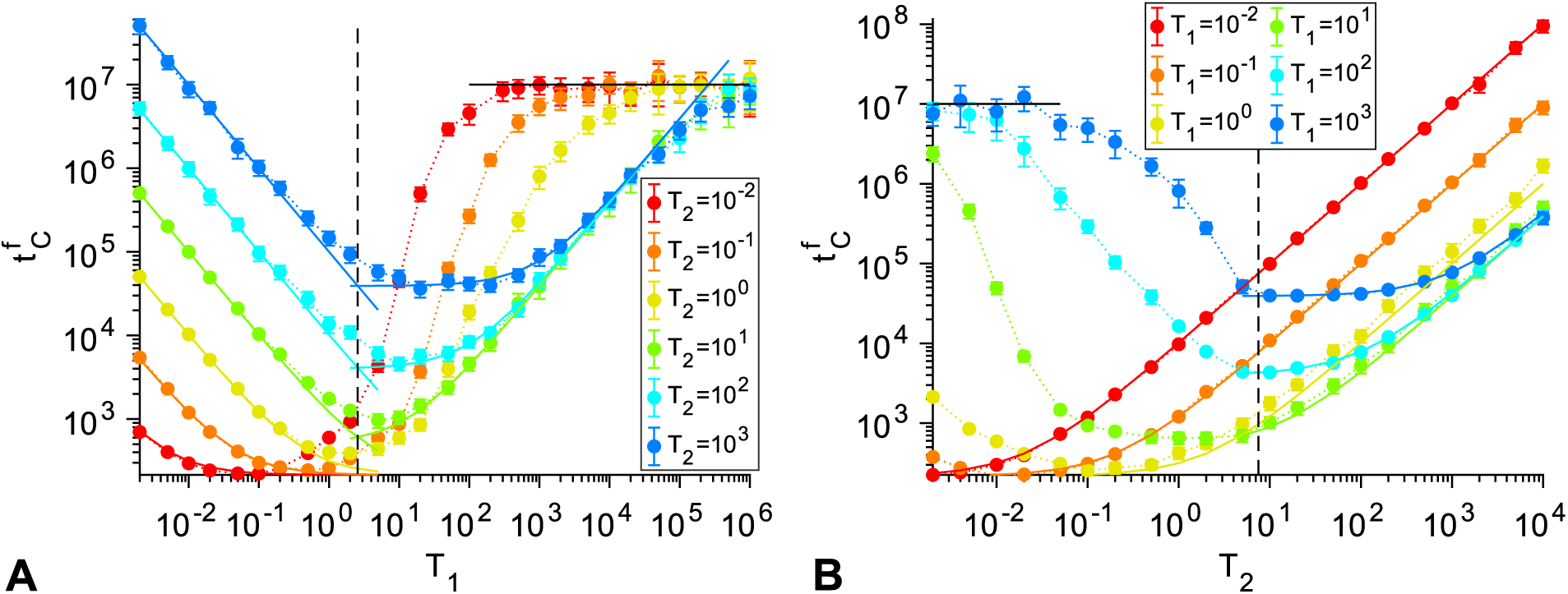
Asymmetric alternations. Fixation time 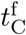 of C individuals in a population subjected to asymmetric alternations of absence and presence of antimicrobial (respective durations: *T*_1_ and *T*_2_). Data points correspond to the average of simulation results, and error bars (sometimes smaller than markers) represent 95% confidence intervals. In both panels, solid lines correspond to our analytical predictions in each regime. Parameter values: *µ*_1_ = 10^−5^, *µ*_2_ = 10^−3^, *δ* = 0.1 and *N* = 10^3^. (A) 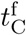 as function of *T*_1_ for different *T*_2_. Dashed line: 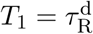 (B) 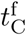 as function of *T*_2_ for different *T*_1_. Dashed line: 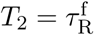

For 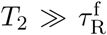, Fig. 5A shows that 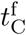 features a striking minimum, which gets higher but wider for longer *T*_2_. This can be fully understood from our analytical predictions. Indeed, when *T*_1_ is varied starting from small values at fixed 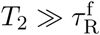, different regimes can be distinguished:

- When 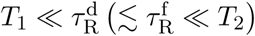 Eq. 3 yields 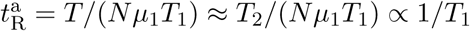
- When 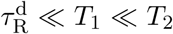 Eq. 3 gives 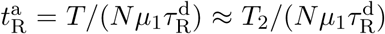, which is independent from *T*_1_.
- As *T*_1_ reaches and exceeds *T*_2_, the law 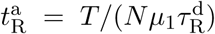 still holds.It yields 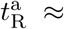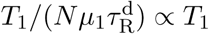 when 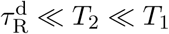.

Hence, the minimum of 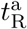 is 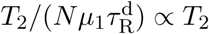 and is attained for 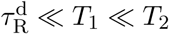 : it gets higher but wider for larger *T*_2_.

In the opposite regime where 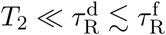, Fig. 5A shows that 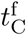 also features a minimum as a function of *T*_1_:

- When 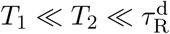, Eq. 3 yields 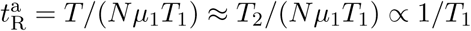
- When 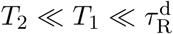, the same law gives 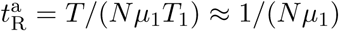, which is independent from *T*_1_.
- When 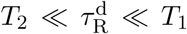, the R lineages that start growing go extinct (see Fig. 4B), and valley crossing then dominates. In Fig 5A, the black horizontal line corresponds to our analytical prediction for valley crossing (see below).

Hence, the minimum of 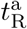 is 1*/*(*Nµ*_1_) and is attained for 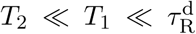 : then, the very first R mutant that appears drives the complete evolution of resistance in the population. For 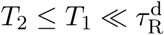, 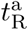 is between once and twice this minimum value.

A similar analysis can be conducted if *T*_2_ is varied at fixed *T*_1_. Fig. 5B shows the corresponding simulation results, together with our analytical predictions from Eqs. 2 and 3. In particular, a minimum is observed in Fig. 5B when varying *T*_2_ for 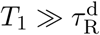

- When 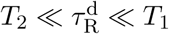, valley crossing dominates.
- When 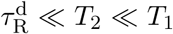, Eq. 3 gives 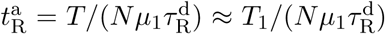, which is independent from *T*_2_.
- As *T*_2_ is further increased, 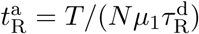 increases, becoming proportional to *T*_2_ when *T*_2_ *≫ T*_1_.

Hence, the minimum of 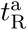 is 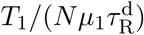 and is attained for 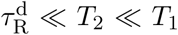. In the opposite case where 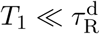, Eq. 3 still gives 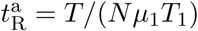 Thus, 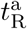 reaches a plateau 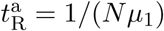 for 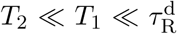, which means that the first R mutant yields the full evolution of resistance (as seen above). Then, 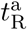 becomes proportional to *T*_2_ for *T*_2_ *≫ T*_1_. Note that valley crossing is always slower than the alternation-driven process when 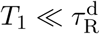 (see below), so no plateau is expected at large *T*_2_ in this case.

In a nutshell, for asymmetric alternations, a striking minimum of the time of full evolution of resistance by a population occurs when both phases have durations of the same order. Interestingly, the minimum generally occurs when the phases of antimicrobial presence are shorter than those of absence, i.e. *T*_2_ *≤ T*_1_ (except if 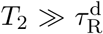). Strikingly, if 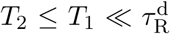, the first R mutant that ever appears in the population ultimately yields the full evolution of resistance, with a timescale of order 1*/*(*Nµ*_1_).

### Comparison to spontaneous fitness valley crossing

**No antimicrobial: Crossing of a symmetric fitness valley**

Let us compare the alternation-driven evolution of resistance to what would happen in the absence of alternations of phases of absence and presence of antimicrobial. If a population composed only of S (sensitive) microorganisms is subjected to a continuous presence of antimicrobial, it will not evolve resistance, because divisions are blocked (see Fig. 1A). Conversely, a population of S microorganisms that is never subjected to antimicrobial can spontaneously evolve resistance. In our model, this will eventually happen. This process is difficult and slow, because of the initial fitness cost of resistance: it requires crossing a fitness valley (see Fig. 1A). Fitness valley crossing has been studied in detail [29, 30, 21, 31, 32], but usually in the case where the final mutant has a higher fitness than the initial organism. In the evolution of antimicrobial resistance, compensatory mutations generally yield microorganisms with antimicrobial-free fitnesses that are similar to, but not higher than those of sensitive microorganisms [3, 10, 4]. Hence, we here extend the known results for fitness valley crossing by constant-size homogeneous asexual populations [21] to “symmetric” fitness valleys, where the final genotype has no selective advantage compared to the initial one.

There are two different ways of crossing a fitness valley. In *sequential fixation*, the first deleterious mutant fixes in the population, and then the second mutant fixes. In *tunneling* [29], the first mutant never fixes in the population, but a lineage of second mutants arises from a minority of first mutants, and fixes. For a given valley, characterized by δ (see Fig. 1A), population size *N* determines which mechanism dominates. Sequential fixation requires the fixation of a deleterious mutant through genetic drift, and dominates for small *N*, when stochasticity is important. Tunneling dominates above a certain *N* [30, 21]. Let us study these two mechanisms for rare mutations *Nµ*_1_ *≪*1.

In sequential fixation, the average time *τ*_SF_ to cross a valley is the sum of those of each step involved [21]. Hence *τ*_SF_ = 1*/*(*Nµ*_1_*p*_SR_) + 1*/*(*Nµ*_2_*p*_RC_), where *p*_SR_ (resp. *p*_RC_) is the fixation probability of a single R (resp. C) individual in a population of size *N* where all other individuals are of type S (resp. R). Fixation probabilities are known in the Moran process (see S1 Appendix). In particular, if *Nδ ≪*1 then *p*_SR_ *≈ p*_RC_ *≈* 1*/N* for our symmetric valley, so *τ*_SF_ *≈* 1*/µ*_1_ + 1*/µ*_2_ (*≈* 1*/µ*_1_ if *µ*_1_ *≪µ*_2_), while if *δ ≪*1 and *Nδ ≫* 1 then *p*_SR_ *≈ δe*^*-Nδ*^ and *p*_RC_ *≈ δ ≫ p*_SR_, so *τ*_SF_ *≈ e*^*Nδ*^/(*Nµ*_1_*δ*).

In tunneling, the key timescale is that of the appearance of a successful first (R) mutant, a first mutant whose lineage will give rise to a second (C) mutant that will fix in the population [21]. Neglecting subsequent second mutation apparition and fixation times, the average tunneling time reads *τ*_T_ *≈* 1*/*(*Nµ*_1_*p*_1_), where *p*_1_ is the probability that a first mutant is successful [21]. Upon each division of a first mutant, the probability of giving rise to a second mutant that will fix is *p* = *µ*_2_*p*_SC_, where *p*_SC_ is the fixation probability of a single C mutant in a population of S individuals. For our symmetric valley, *p*_SC_ = 1*/N*, so *p* = *µ*_2_*/N*. In the neutral case *δ* = 0, Ref. [21] demonstrated that the first-mutant lineages that survive for at least 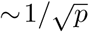 generations, and reach a size 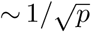, are very likely to be successful, and fully determine the rate at which successful first mutants are produced. Since the lineage of each new first mutant has a probability 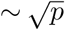 of surviving for at least 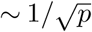 generations [21], the probability that a first mutant is successful is 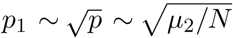. If *δ >* 0, a first mutant remains effectively neutral if its lineage size is smaller than 1*/δ* [21]. Hence, if 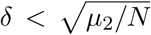, 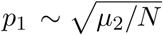 still holds. (This requires Nµ2≫ 1, otherwise the first mutant fixes before its lineage reaches a size 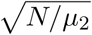 Finally, if 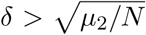, the lineage of a first mutant will reach a size at most *∼* 1*/δ*, with a probability *∼ δ* and a lifetime *∼* 1*/δ* [21], *yielding p*_1_ *∼ µ*_2_*/*(*Nδ*).

Given the substantial cost of resistance mutations (*δ ∼* 0.1 [10, 4]) and the low compensatory mutation rates (in bacteria *µ*_2_ *∼* 10^−8^ [4]), let us henceforth focus on the case where 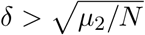 (which is appropriate for all *N ≥* 1 with the values mentioned). Then *τ*_T_ *≈* 1*/*(*Nµ*_1_*p*_1_) *≈δ/*(*µ*_1_*µ*_2_), and two extreme cases can be distinguished:

*(A) Nδ ≪*1 (effectively neutral regime): Then, *τ*_SF_ *≈* 1*/µ*_1_ (for *µ*_1_ *≪µ*_2_) and *τ*_T_ *≈ δ/*(*µ*_1_*µ*_2_). Given the orders of magnitude above, generally *δ > µ*_2_ in resistance evolution. Hence, sequential fixation is fastest, and the valley crossing time *τ*_V_ reads:

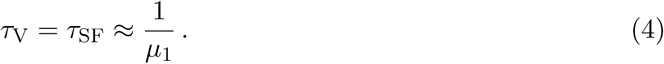

(B) *δ* ≪1 and *Nδ ≫* 1: Then,

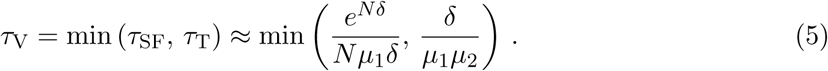

The transition from sequential fixation to tunneling [21] occurs when *Nδe*^*-Nδ*^ = *µ*_2_*/δ*.

We have focused on the rare mutation regime *Nµ*_1_ *≪*1. If mutations are more frequent, the first successful lineage of R mutants that appears may not be the one that eventually fixes, so the valley-crossing time becomes shorter [21].

In Fig. 3B, the black simulation data points were obtained without any antimicrobial. The population then evolves resistance by valley crossing. The black curves correspond to our analytical predictions in Eq. 4 for *N ≪* 1 */δ* and in Eq. 5 for *N ≫* 1*/δ*. In the latter regime, the transition from sequential fixation to tunneling occurs at *N ≈* 65 for the parameters of Fig. 3B. The agreement between simulation results and analytical predictions is excellent, with no adjustable parameter.

### Alternation-driven process vs. valley-crossing process

Now that we have studied the spontaneous crossing of a symmetric fitness valley without any antimicrobial, let us come back to our periodic alternations of phases of absence and presence of antimicrobial. Resistance can then evolve by two distinct mechanisms, namely the alternation-driven process and the spontaneous valley-crossing process. It is important to compare the associated timescales, in order to assess which process will happen faster and dominate. This will shed light on the acceleration of resistance evolution by the alternations. For generality, we consider asymmetric alternations.

With alternations, spontaneous valley crossing can still happen, but new R lineages cannot appear with antimicrobial, because S individuals cannot divide (see Fig. 1A). Since the appearance of a successful R mutant is usually the longest step of valley crossing (see above), the average valley crossing time 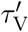 with alternations will be longer by a factor *T/T*_1_ than that with-out antimicrobial (*τ*_V_), if more than one antimicrobial-free phase is needed to cross the valley, if *T*_1_ *≪τ*_V_. Eqs. 4 and 5 then yield

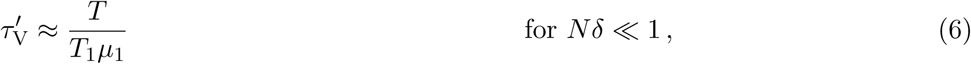

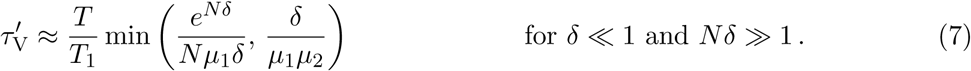

Conversely, if *T*_1_ *≫ τ*_V_, valley crossing generally happens within the first antimicrobial-free phase. Hence, the average valley crossing time *τ*_V_ is given by Eqs. 4 and 5. (Recall that the process is assumed to begin with an antimicrobial-free phase.)

We can now compare the timescales of the valley-crossing process to those of the alternation-driven process. For simplicity, let us assume that the dominant timescale in the latter process is the time 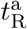 it takes to first observe a R organism in the presence of antimicrobial, i.e. 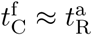 (see Eq. 2). This is the case in a large and relevant range of parameters, especially if *µ*_1_ ≪ *µ*_2_, as discussed above. Note also that the final step of fixation of the successful C lineage, which can become long in large populations (*∼ N*, see S1 Appendix), is the same in the alternation-driven process and in the valley-crossing process, so it does not enter the comparison. The expression of 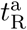 in Eq. 3 should thus be compared to the valley crossing time. If *T*_1_ *≫ τ*_V_, valley crossing happens before any alternation, and is thus the relevant process, with time *τ*_V_ given by Eqs. 4 and 5. Let us now conduct our comparison of 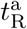 and 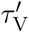 for *T*_1_ ≪ *τ*_V_, where Eqs. 6 and 7 hold.

(A) If 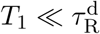 (recall that 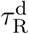 is the average lifetime of an R lineage without antimicrobial, before it goes extinct): The alternation-driven process, with timescale 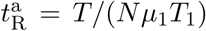 (see Eq. 3), dominates. Indeed, if *Nδ ≪*1, 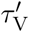 is given by Eq. 6, so for all *N >* 1, 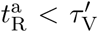 And if *Nδ ≫* 1 and *δ ≪*1, Eq. 7 yields 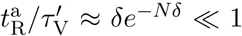 in the sequential fixation regime, and 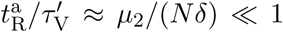 in the tunneling regime. Hence, if 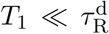, alternations of absence and presence of antimicrobial strongly accelerate resistance evolution. For instance, in Fig. 3A, for *N* = 100 and 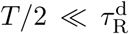, the alternation-driven process takes 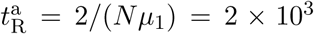 generations, while valley crossing takes *τ*_V_ = *δ/*(*µ*_1_*µ*_2_) = 10^7^ generations without antimicrobial: alternations yield a speedup of 4 orders of magnitude. The speedup is even stronger for larger populations.

(B) If 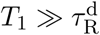 : Then 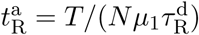 (see Eq. 3). If *Nδ ≪*1, valley crossing by sequential fixation is the dominant process. Indeed, Eq. 6 yields 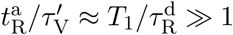 and *δ ≪*1, Eq. 7 yields 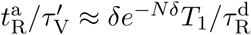 in the sequential fixation regime, and 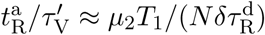 in the tunneling regime. A transition from the alternation-driven process to valley crossing occurs when these ratios reach 1. Qualitatively, if *N* is large enough and/or if *T*_1_ is short enough, the alternation-driven process dominates.

For example, in Fig. 5A, parameters are such that the dominant mechanism of valley crossing is tunneling, so 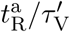 reaches 1 for 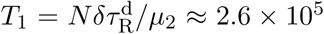 generations. This transition to the valley-crossing plateau is indeed observed for the curves with large enough *T*_2_. (Recall that if 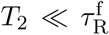, extinction events occur when 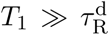, see Fig. 4B.) The black horizontal lines in Figs. 5A and 5B correspond to our analytical prediction in Eq. 7, giving 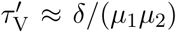 if 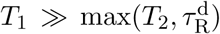. Similarly, in Fig. 3A, horizontal solid lines at large *T* correspond to the valley crossing times in Eqs. 6 or 7, depending on *N*. In Fig. 3B, in the regime of small *N* and large *T*, resistance evolution is achieved by tunneling-type valley crossing, yielding a plateau in the neutral regime *N ≪*1*/δ* (see Eq. 6, plotted as a horizontal purple line) and an exponential increase for intermediate *N* (see Eq. 7). For larger *N*, we observe a *T*-dependent transition to the alternation-driven process, which can be fully understood using the ratio 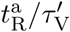 (see above).

## Discussion

### Main conclusions

Because of the generic initial fitness cost of resistance mutations, alternations of phases of absence and presence of antimicrobial induce a dramatic time variability of the relevant adaptive landscape, which alternates back and forth from a fitness valley to an ascending landscape. Using a generic and minimal theoretical model which retains the key biological ingredients, we have shed light on the quantitative implications of these time-varying patterns of selection on the time it takes for resistance to fully evolve *de novo* in a homogeneous microbial population of fixed size. Combining analytical approaches and simulations, we showed how resistance evolution can be driven by periodic alternations of phases of absence and presence of an antimicrobial that stops growth. We compared this alternation-driven process to the spontaneous valley-crossing process. We quantified how the time necessary for resistant-compensated microorganisms to take over a microbial population depends on the alternation period.

We found that fast alternations strongly accelerate the evolution of resistance, reaching a plateau once the half-period *T/*2 becomes shorter than the average lifetime 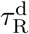 before extinction of a resistant lineage without antimicrobial. Strikingly, in this case, the very first resistant mutant that appears ultimately leads to full resistance of the population. Conversely, when 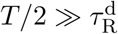, the time needed for resistance to evolve increases linearly with *T*, until it reaches the spontaneous valley-crossing time, which constitutes an upper bound. Our full-fledged stochastic model allowed us to investigate the impact of population size *N*. We showed that the acceleration of resistance evolution is stronger for larger populations, eventually reaching a plateau in the deterministic limit. The valley-crossing plateau is reached in the opposite limit of small populations, most strikingly if the first resistance mutation is effectively neutral. Over a large range of intermediate parameters, the time needed for the population to fully evolve resistance scales as *T/N*. These results are summed up in Fig. 6A.

**Figure 6:**
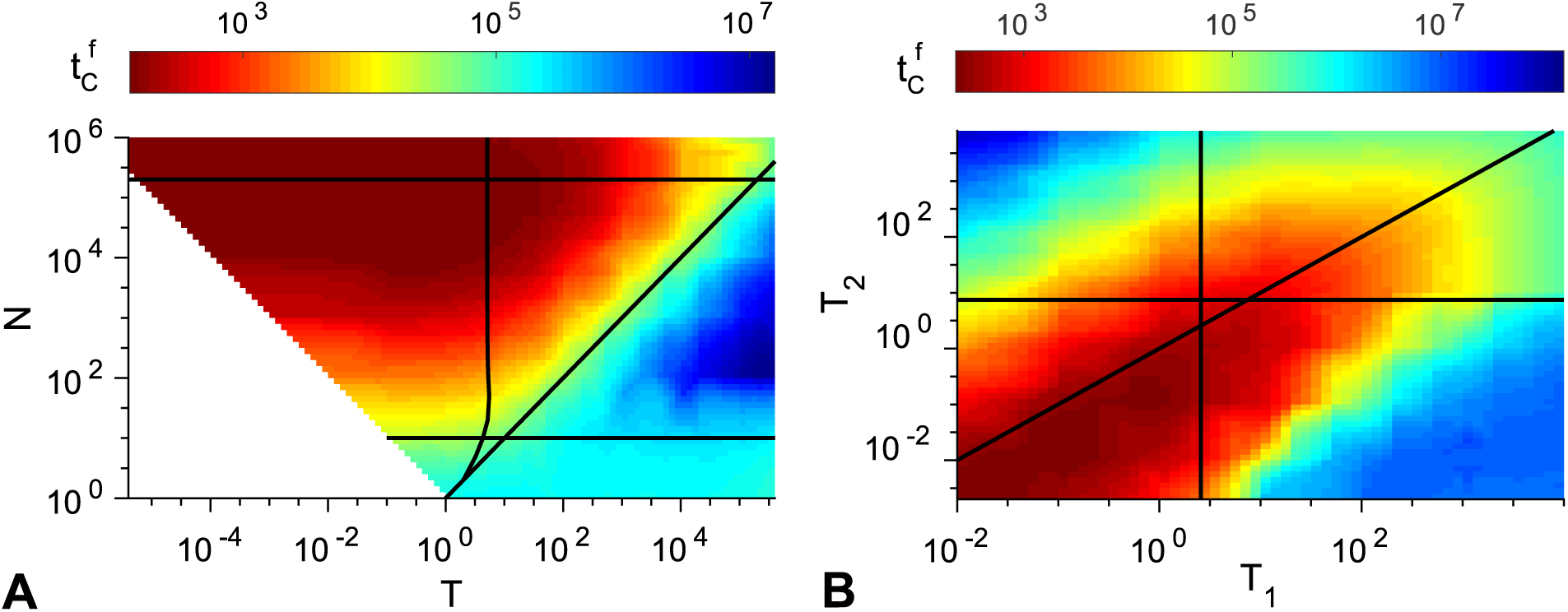
Heatmaps. Fixation time 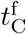 of C individuals in a population of size *N* subjected to periodic alternations of absence and presence of antimicrobial. Simulation data plotted in Figs. 3A and 5A are linearly interpolated. Parameter values: *µ*_1_ = 10^−5^, *µ*_2_ = 10^−3^, *δ* = 0.1. (A) Symmetric alternations: 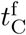 as function of the period *T* and the population size *N*. Top horizontal line: deterministic regime limit *N* = 1*/µ*_1_. Bottom horizontal line: neutral regime limit *N* = 1*/δ*. Quasi-vertical curve: 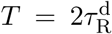. Diagonal line: *T* = *N*. (B) Asymmetric alternations: 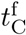 as function of the durations *T*_1_ and *T*_2_ of the phases of absence and presence of antimicrobial. Vertical line: 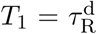. Horizontal line: 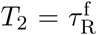. Diagonal line: *T*_1_ = *T*_2_. Here *N* = 10^3^, so the first resistant mutant appears after an average time *T/*(*Nµ*_1_*T*_1_) = 10^2^ *T/T*_1_.

For asymmetric alternations, featuring different durations *T*_1_ and *T*_2_ of the phases of absence and presence of antimicrobial, we have shed light on the existence of a minimum of the time taken by the population to fully evolve resistance. This striking minimum occurs when both phases have durations of the same order, generally with *T*_1_ *≤ T*_2_. Moreover, the minimum value reached for the time of resistance evolution becomes smaller for shorter alternation periods. When they are shorter than 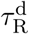, the full evolution of resistance is triggered by the first R mutant that appears in the population. These results are summed up in Fig. 6B.

### Context and perspectives

Our approach is complementary to previous studies providing a detailed modeling of specific treatments [33, 34, 20, 35, 36, 37, 38]. Indeed, the majority of them [33, 39, 34, 20, 35, 36, 40] neglect stochastic effects, while they can have a crucial evolutionary impact [41, 25, 42]. The deterministic approach is appropriate if the number *N* of competing microbes satisfies *Nµ*_1_ *≫* 1, where *µ*_1_ is the mutation rate [43, 41]. Such large sizes can be reached in some established infections [22], but microbial populations go through very small bottleneck sizes (sometimes *N ∼* 1 *-* 10 [44, 45, 46]) when an infection is transmitted. Moreover, established microbial populations are structured, even within a single patient [47, 48], and competition is local, which decreases the effective value of *N*. Some previous studies did take stochasticity into account, but several did not include compensation of the cost of resistance [49, 50], while others made specific epidemiological assumptions [37].

Our model assumes that the size of the microbial population remains constant. While this is realistic in some controlled experimental setups, e.g. chemostats [26], microbial populations involved in infections tend to grow, starting from a small transmission bottleneck, and the aim of the antimicrobial treatment is to make them decrease in size and eventually go extinct. In the case of biostatic antimicrobials, which prevent bacteria from growing (as in our model), populations can go extinct due to spontaneous and immune system-induced death. Our model with constant population size should however be qualitatively relevant at the beginning and middle stages of a treatment (i.e. sufficiently after transmission and before extinction). Constant population sizes facilitate analytical calculations, and allowed us to fully quantify the impact of a periodic presence of antimicrobial on resistance evolution, but it will be very interesting to extend our work to variable population sizes [51, 52, 16, 53, 54]. Another interesting extension would be to incorporate spatial structure [32, 55, 56, 57] and environment heterogeneity, in particular drug concentration gradients. Indeed, static gradients can strongly accelerate resistance evolution [58, 59, 60, 61], and one may ask how this effect combines with the temporal alternation-driven one investigated here.

### Implications for clinical and experimental situations

The situation where *T*_1_ and *T*_2_ are of the same order and shorter than 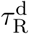 yields a dramatic acceleration of the evolution of resistance, and is unfortunately clinically realistic. Indeed, a goal in treatment design is that the serum concentration of antimicrobial exceeds the MIC for at least 40 to 50% of the time [62], which implies that many actual treatments may involve the alternations that most favor resistance evolution according to our results [62, 63, 20]. Besides, for bacteria dividing on a timescale of about an hour, 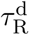 is a few hours. Since antimicrobial is often taken every 8 to 12 hours in treatments by the oral route, having *T*_1_ and *T*_2_ of similar order as 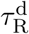 is realistic.

In the worst case where *T*_1_ and *T*_2_ are of the same order and shorter than 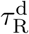, full *de novo* resistance evolution results from the apparition of the very first R mutant, which takes *T/*(*Nµ*_1_*T*_1_). Under the conservative assumption that only one resistance mutation is accessible, taking *µ*_1_ *∼* 10^−10^, which is the typical mutation probability per nucleotide and per generation in *Escherichia coli* bacteria [64], we find that this duration is less than a day (*∼* 10*-*20 generations) for *N ∼* 10^9^, and a few days for *N ∼* 10^8^, numbers which can be reached in infections [22]. For such large populations, the fixation of the C (compensated) mutant will take more time, but once R is fixed (which takes *∼* 1 day after the appearance of the first R mutant), C will fix even if the treatment is stopped. This is due to the large number of compensatory mutations, which yields a much higher effective mutation rate toward compensation than toward reversion to sensitivity [7, 8, 22]. In addition, many mutations to resistance are often accessible, yielding higher effective *µ*_1_, *µ*_1_ *∼* 10^−8^ for rifampicin resistance in some wild isolates of *E. coli* [9], meaning that smaller populations can also quickly become resistant in the presence of alternations. Recall that we are only considering *de novo* resistance evolution, without pre-existent resistant mutants, or other possible sources of resistance, such as horizontal gene transfer, which would further accelerate resistance acquisition.

In summary, an antimicrobial concentration that drops below the MIC between each intake can dramatically favor *de novo* resistance evolution. More specifically, we showed that the worst case occurs when *T*_1_ *≤ T*_2_, which would be the case if the antimicrobial concentration drops below the MIC relatively briefly before each new intake. Our results thus emphasize how important it is to control for such apparently innocuous cases, and constitute a striking argument in favor of the development of extended-release antimicrobial formulations [65].

While the parameter range that strongly accelerates resistance evolution should preferably be avoided in clinical situations, it could be tested and harnessed in evolution experiments. Again, these parameters are experimentally accessible. Periodic variations of antimicrobial concentrations are already used experimentally, in particular in morbidostat experiments [66, 67], where the population size is kept almost constant (as in chemostats more generally), which matches our model. In Ref. [66], a dramatic and reproducible evolution of resistance was observed in *∼* 20 days when periodically adjusting the drug concentration to constantly challenge *E. coli* bacteria. Given our results, it would be interesting to test whether resistance evolution could be made even faster by adding drug in a chemostat with a fixed periodicity satisfying 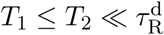.

## Acknowledgments

We thank Claude Loverdo, David J. Schwab and Raphaėl Voituriez for stimulating discussions. AFB also acknowledges the KITP Program on Evolution of Drug Resistance (KITP, Santa Barbara, CA, 2014), which was supported in part by the National Science Foundation under Grant NSF PHY 17-48958.

## Supporting Information

**Table S1:**
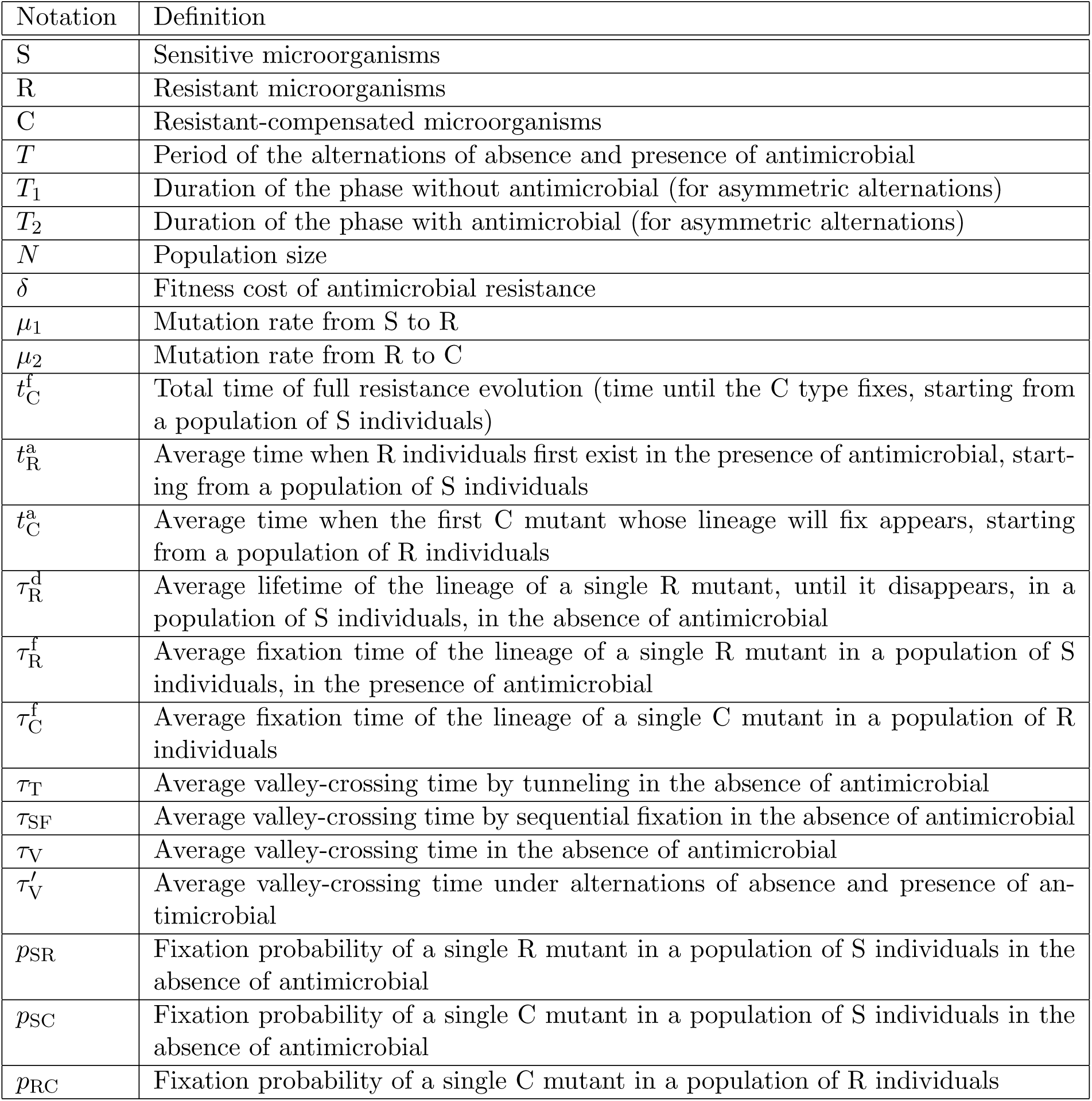
Notations. This table lists the different notations introduced in the main text and their meaning.

**Figure S1:**
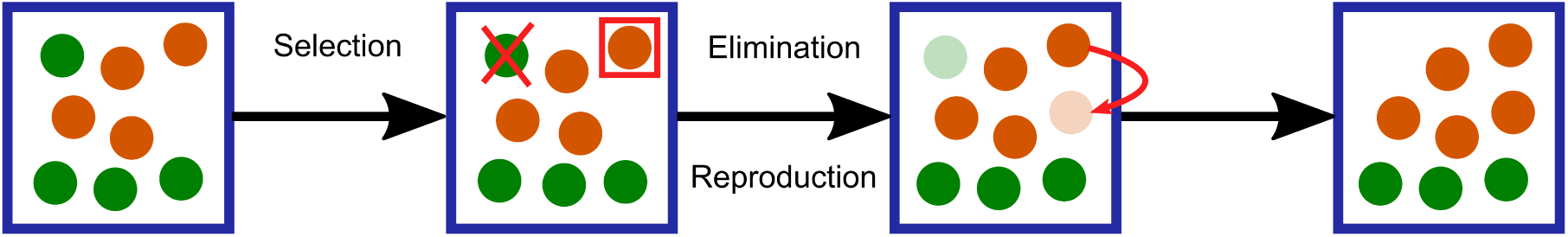
Sketch of the Moran process. One step of the Moran process is represented in a population with 8 individuals of 2 different types (different colors).

**Figure S2:**
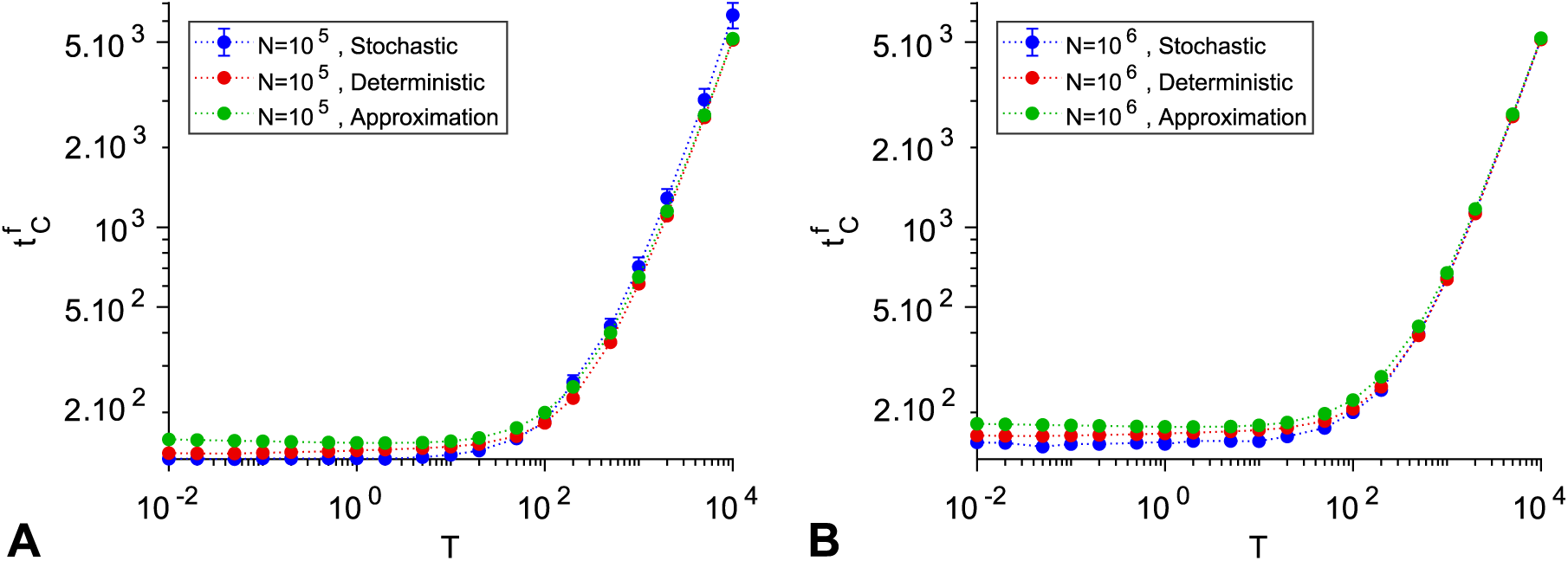
Large populations: stochastic model vs. deterministic model. The total time *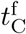* of full resistance evolution is plotted versus the period *T* of alternations of absence and presence of antimicrobial, in the case of symmetric alternations. Results from simulations of the stochastic model (see Fig. 3A), numerical resolution of the deterministic model, and an analytical approximation of the deterministic solution (Eqs. S60-S61), are represented for *N* = 10^5^ (A) and *N* = 10^6^ (B). Parameter values: *µ*_1_ = 10^−5^, *µ*_2_ = 10^−3^, and *δ* = 0.1.

**Figure S3:**
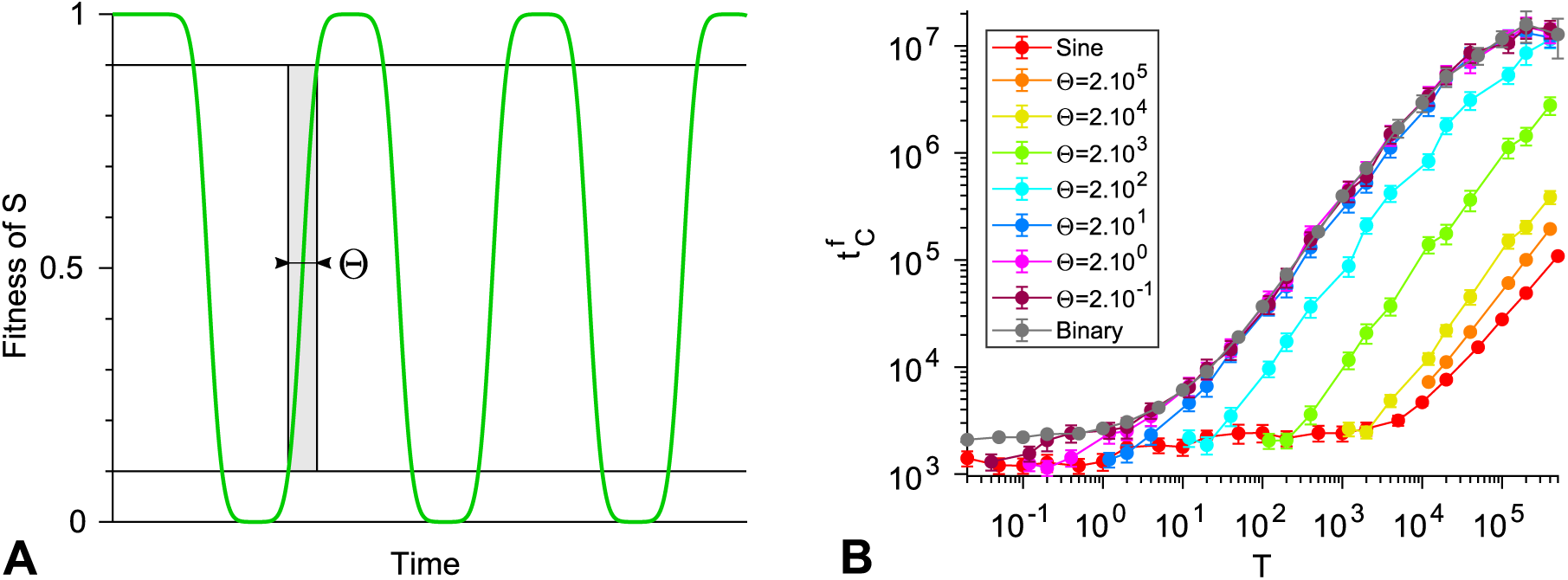
Robustness of the binary antimicrobial action model. (A) Smooth and periodic fitness versus time relationship considered: Θ denotes the rise time. (B) Total time 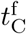 of full resistance evolution versus the period *T* for smooth alternations with different values of Θ, and for the binary model. Data points correspond to the average of simulation results, and error bars (often smaller than markers) represent 95% confidence intervals. Parameter values: *µ*_1_ = 10^−5^, *µ*_2_ = 10^−3^, *δ* = 0.1, and *N* = 100.

## S1 Appendix

### 1. Fixation probabilities and fixation times in the Moran process

Here, we discuss in detail the fixation probabilities and mean fixation times in the Moran process, which are used throughout the main text. These quantities are already known [25, 28], but we present a derivation for the sake of pedagogy and completeness. Our derivation is based on the general formalism of first passage times, and gives the same results as those obtained in the literature, often using other methods [25, 28]. Next, we use the general expressions obtained to express the various fixation probabilities and fixation times used in the main text.

#### 1.1 The Moran process

The Moran model [24, 25] is a simple stochastic process used to describe the evolution of the composition of asexual populations of finite and constant size. It allows to incorporate variety increasing processes such as mutation and variety-reducing processes such as natural selection.

In the Moran model, at each time step, an individual is chosen at random to reproduce and another one is chosen to die (see Figure S1). Hence, the total number of individuals in the population stays constant. Note that we will consider that the same individual can be selected to reproduce and die at the same step. Natural selection can be introduced by choosing the individual that reproduces with a probability proportional to its fitness. To implement mutations upon division, one can allow the offspring to switch type with a certain probability at each step. When a mutant arises within the Moran model, its lineage can either disappear or fix in the population, i.e. take over the whole population. The outcome is not fully determined by fitness differences as in a deterministic case, but also by stochastic fluctuations, also known as genetic drift. Here, we focus on the evolution of population composition under genetic drift and selection alone. In the rare mutation regime, these processes are much faster than the time between the occurrence of two mutations, so mutation can be neglected during the process of fixation of one type. The Moran model allows to compute explicit expressions for quantities such as fixation probabilities and fixation times [25, 68] (see below).

Let us consider a population of *N* individuals of two types A and B, which have fitnesses *f*_*A*_ and *f*_*B*_, respectively. We denote the number of A individuals by *j*. Thus *N - j* represents the number of B individuals. Let us study the evolution of *j* at one step of the Moran process (for an example, see Figure S1). The transition probabilities associated to the Moran process read [25]:

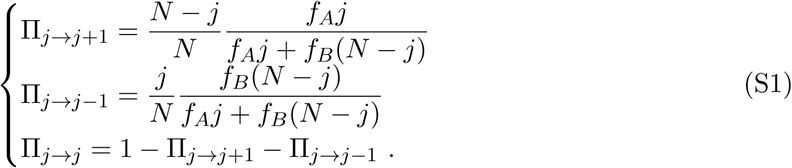

The Moran process is a discrete-time Markov process, since the probabilities of states *j* after one step only depend upon the present value of *j* [69]. Let us take the limit of continuous time and write the master equation 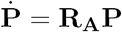 giving the probability of being at state *j* at time *t*:

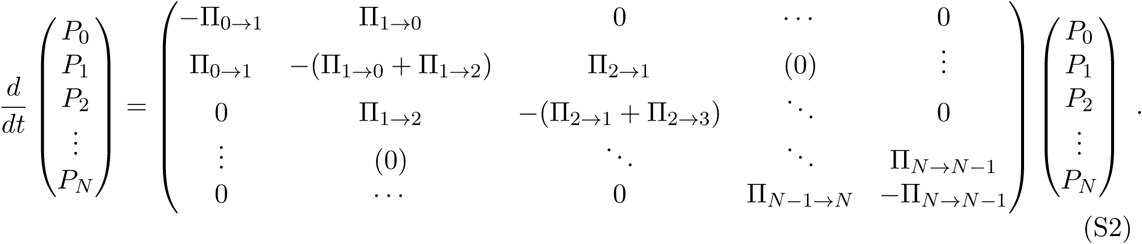

This Markov chain has two absorbing states, namely *j* = 0 and *j* = *N*, which correspond to the fixation of B and A individuals, respectively. Once these states are reached, no more changes can occur, in the absence of mutation. It follows that all the components of the first and the last columns of **R_A_** equal to 0 (see Eq. S2), so **R_A_** is not invertible. In the following, we will denote by 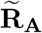 the reduced transition rate matrix in which the rows and the columns corresponding to the absorbing states (*j* = 0, *j* = *N*) are removed, and by 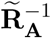 its inverse. Let us note that **R_A_** is a tridiagonal matrix, which allows for major simplifications of analytical calculations [70]. Note that in order to obtain the transition rate matrix associated to B individuals, one just needs to apply the reversal *j ↔ N - j*. This corresponds to using the matrix **RB** = **JR_A_J** where **J** is the anti-identity matrix. For instance, in 2 dimensions,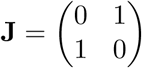

#### 1.2 General fixation probabilities and fixation times

##### Definitions

The fixation probability 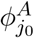 represents the probability that A individuals finally succeed and take over the population, starting from *j* = *j*_0_ individuals of type A. In particular, 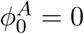 and 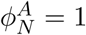. Similarly, 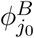 is the fixation probability of the B individuals, still starting from *j* = *j*_0_ individuals of type A.

Mean fixation times are the mean times to reach one of the absorbing states. The unconditional fixation time *t*_*j*_0 is the average time until fixation in either *j* = 0 or *j* = *N*, when starting from a number *j* = *j*_0_ of A individuals. The conditional fixation time 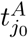 corresponds to the average time until fixation in *j* = *N*, when starting from *j*_0_, provided that type A fixes. Note that in what follows, we will express the fixation times in numbers of steps of the Moran process. Conversion to real times can then be performed by dividing the number of Moran steps by *N*.

In the following, we present a derivation of the fixation probabilities and of the fixation times in the Moran process [25, 28] that uses the general formalism of mean first passage times [71].

##### Fixation probabilities

Assuming that at *t* = 0, the system is at state *j* = *j*_0_, let us focus on the fixation probability 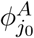 of the A type in the population. The stochastic process stops at the time 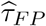 when *j* fixes, i.e. first reaches one of the absorbing states *{j* = 0, *j* = *N }*. Hence, integrating over all values of 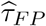, under the condition that fixation finally occurs in *j* = *N*, yields

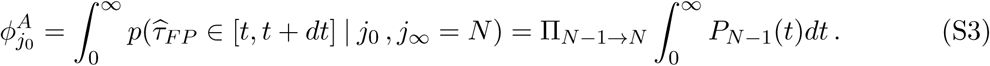

In the last expression, we have taken advantage of the fact that the only way to fix in *j* = *N* between *t* and *t* + *dt* is to be in state *j* = *N -* 1 at time *t* and then to transition from *N -* 1 to *N* (see Eq. S2). We have thus introduced the probability *P*_*N-*1_(*t*) of being in state *j* = *N -* 1 at time *t*, starting in state *j* = *j*_0_ at time 0. More generally, the probability *P*_*i*_(*t*) can be considered.

Integrating the Master equation Eq. S2 to determine *P*_*i*_(*t*), with the initial condition *P*_*i*_(0) = *δ*_*i* *j*_0, yields

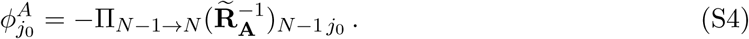

A similar reasoning gives the fixation probability 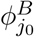 of the B type, still starting from *j*_0_ individuals of type A and *N - j*_0_ individuals of type B:

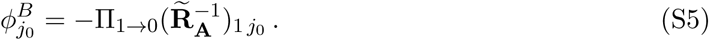

These two probabilities satisfy 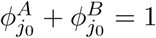 since there are 2 absorbing states in the process.

##### Mean fixation times

Let us now focus on the mean fixation times, still assuming that at *t* = 0, the system is at state *j* = *j*_0_. The probability that fixation in one of the absorbing states *{j* = 0, *j* = *N }* occurs between *t* and *t* + *dt* reads:

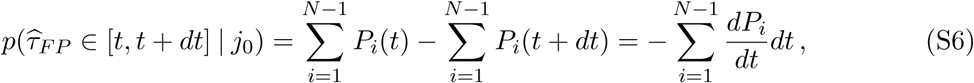

where, as above, *P*_*i*_(*t*) represents the probability of being in state *i* at time *t* starting in *j*_0_ at time 0 (note that the initial condition *j*_0_ is omitted for brevity). Thus, the unconditional fixation time can be expressed as:

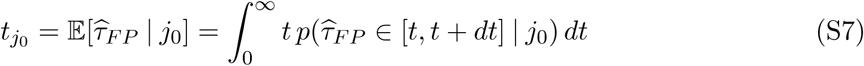

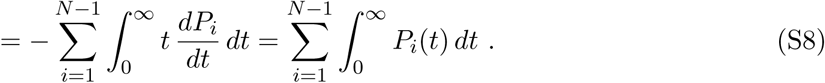

Integrating the Master equation Eq. S2 to determine *P*_*i*_(*t*), with the initial condition *P*_*i*_(0) = *δ*_*i* *j*_0, gives

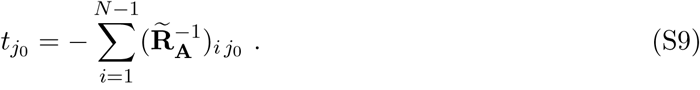

To express the conditional fixation time 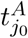 of type A, starting from *j*_0_ A individuals, we need to take into account the condition that fixation finally occurs in state *j* = *N* :

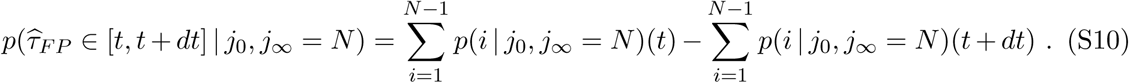

The Bayes relation [72] gives:

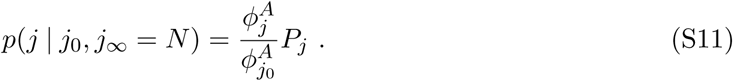

By using the same method as for the unconditional fixation time, one obtains:

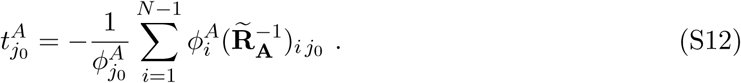

Similarly, the conditional fixation time of the B type, starting from *j*_0_ A individuals, reads:

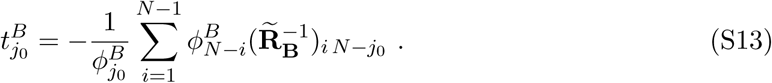

It is straightforward to verify that Eqs. S9, S12 and S13 are linked by the relation:

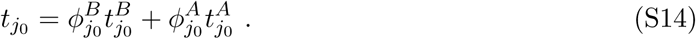

##### Neutral drift

Let us first consider the case without selection *f*_*A*_ = *f*_*B*_. In this case, the Moran process can be seen as a non-biased random walk, since individuals of both types are equally likely to be picked for reproduction and suppression. Fixation eventually happens due to fluctuations. This process, called neutral drift [73] corresponds to diffusion in physics. The transition rates of the system (S1) simplify as follows:

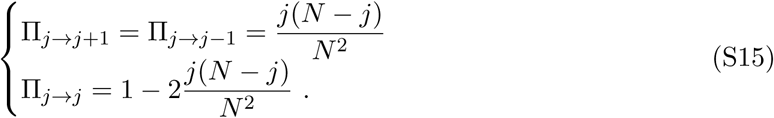

Note that here, *j* can denote the number of A or B individuals indifferently. Indeed, the symmetry *j ↔ N - j* entails **R_A_** = **R_B_** = **R**, and the transition rate matrix is centrosymmetric, i.e. **R** = **JRJ** [74, 75]. For consistency, we will continue to call *j* the number of A individuals.

The fixation probability 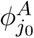 can be obtained from Eq. S4. It involves elements of the inverse of the transition rate matrix. Solving 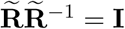 where **I** is the identity matrix, gives

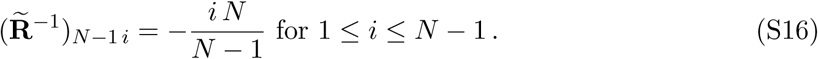

Hence,

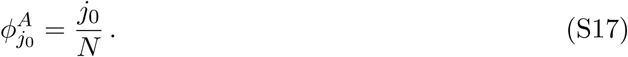

Taking advantage of the centrosymmetry of R (see above), a property which transfers to 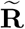 and 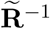, and entails 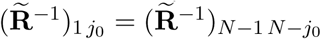, we can apply Eq. S5, yielding

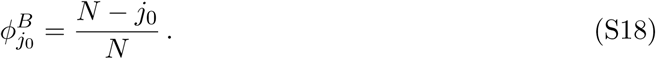

Note that 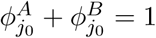, as expected.

Let us now express the fixation times, focusing on the fate of a single mutant of type B, which corresponds to *j*_0_ = *N -* 1. To compute the unconditional fixation time *t*_*N-*1_, we again need elements of the inverse of the transition rate matrix (see Eq. S9), which are given by

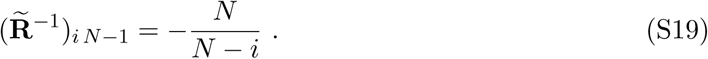

Using Eqs. S9 and S19, we obtain:

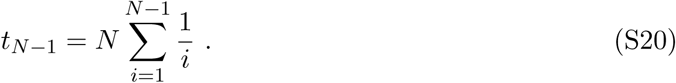

Similarly, using Eqs. S12, S17 and S19, we obtain the conditional fixation time of type A:

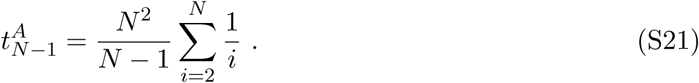

Finally, using Eqs. S13, S17 and S19, and making use of the centrosymmetry of 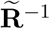 (see above), yields the conditional fixation time of type B:

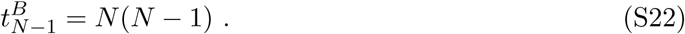

##### Selection

Let us now study the more general case involving selection. For this, let us consider two types A and B having different fitnesses *f*_*A*_ and *f*_*B*_, and let us introduce *γ* = *f*_*A*_*/f*_*B*_. Note that with selection, the transition rate matrices **R_A_** and **R_B_** = **JR_A_J** are different. In order to compute the fixation probability 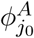, we need some elements of the inverse of the transition rate matrix 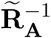, which are given by:

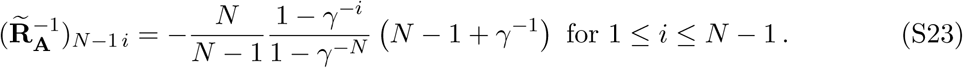

Then, using the previous result and Eq. S4, one obtains:

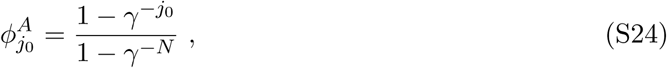

and 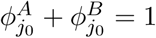 yields:

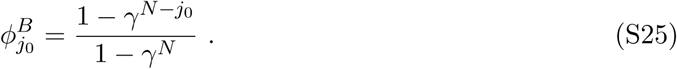

Let us now turn to the fixation times. According to Eq. S9, we need to compute other elements of the inverse of the transition rate matrix 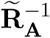 Those satisfy:

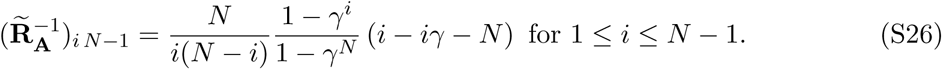

Using Eqs. S9 and S26, the unconditional fixation time reads:

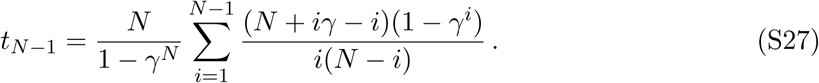

To compute the conditional fixation time 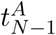, we substitute Eqs. S24 and S26 in Eq. S12, obtaining:

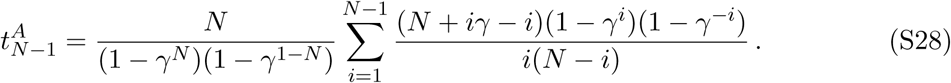

A similar reasoning can be used to obtain the conditional fixation time 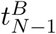 starting from Eq. S13. In order to express the required 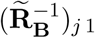, we combine the relation 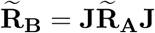, which implies 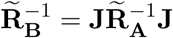 together with Eq. S26, and obtain

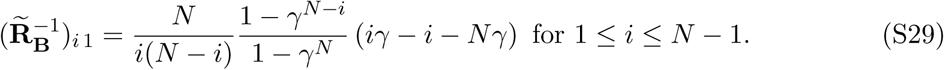

This finally yields

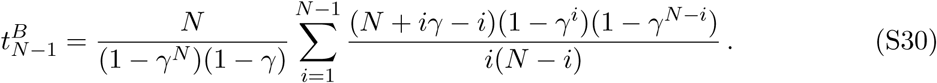

#### 1.3 Fixation probabilities and fixation times used in the main text

Let us now make an explicit link between the general expressions obtained above and the fixation probabilities and fixation times used in the main text.

##### Fixation probabilities

First, in the main text, *p*_SR_ represents the probability that a single resistant (R) mutant fixes without antimicrobial in a population of size *N* where all other individuals are of type S. Without antimicrobial, *f*_S_ = 1 and *f*_R_ = 1 *- δ*. Considering S as type A and R as type B, we have *γ* = *f*_S_*/f*_R_ = 1*/*(1 *- δ*), and our initial condition is *j*_0_ = *N -* 1. Hence, Eq. S25 yields

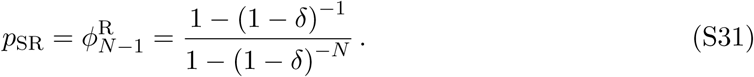

In particular, in the effectively neutral case where *δ ≪*1 and *Nδ ≪*1, it yields

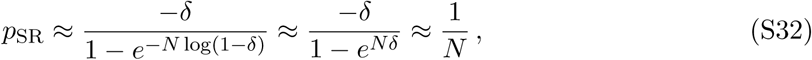

i.e. we recover the result of the neutral case *δ* = 0 (see Eq. S17). Conversely, in the regime where *δ ≪*1 and *Nδ ≫* 1, Eq. S31 yields

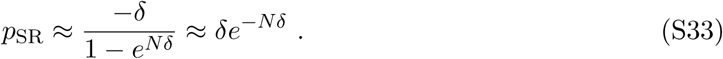

Second, *p*_RC_ denotes the fixation probability of a single C individual in a population of size *N* where all other individuals are of type R. Independently of antimicrobial presence, *f*_R_ = 1 *- δ* and *f*_C_ = 1. Considering R as type A and C as type B, we have *γ* = *f*_R_*/f*_C_ = 1 *- δ*, and our initial condition is *j*_0_ = *N -* 1. Hence, Eq. S25 yields

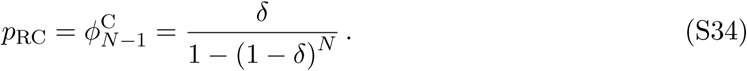

In particular, in the effectively neutral case where *δ ≪*1 and *Nδ ≪*1, it yields

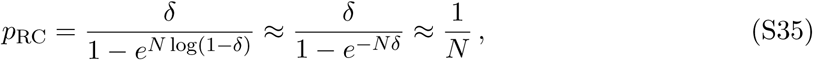

i.e. we again recover the result of the neutral case *δ* = 0 (see Eq. S17). Conversely, in the regime where *δ ≪*1 and *Nδ ≫* 1, Eq. S34 yields

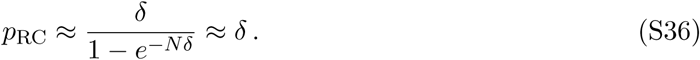

Finally, *p*_SC_ denotes the fixation probability of a single C mutant in a population of S individuals, without antimicrobial. In this case, *f*_S_ = *f*_C_ = 1, so we are in the neutral case, and Eq. S17 yields *p*_SC_ = 1*/N*.

##### Fixation times

First, 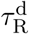 denotes the average time it takes for the lineage of a single R mutant to disappear in the absence of antimicrobial. Hence, it is equal to the fixation time of the S type in a population that initially contains *N -* 1 individuals of type S and 1 individual of type R. Considering S as type A and R as type B, we have *γ* = *f*_S_*/f*_R_ = 1*/*(1 *- δ*) without antimicrobial, and our initial condition is *j*_0_ = *N -* 1, so 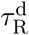 is equal to 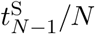 (see Eq. S28). Recall that 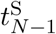 needs to be divided by the population size *N* because we expressed it in numbers of steps of the Moran process, while 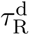 has to be expressed in numbers of generations. While the general formula Eq. S28 is rather complex, in the neutral case *δ* = 0, it reduces to the much simpler expression in Eq. S21, which yields 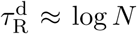 for *N ≫* 1. For 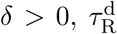 is shorter than in the neutral case, because the R mutants are out-competed by S individuals.Note that a good approximation to the exact formula in Eq. S28 can be obtained within the diffusion approach [25] (see the Fokker-Planck equation below).

Second, 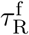 denotes the average time needed for the R mutants take over with antimicrobial, starting from one R mutant and *N -* 1 S individuals. Considering S as type A and R as type B, we have *γ* = *f*_S_*/f*_R_ = 0 with antimicrobial, and our initial condition is *j*_0_ = *N -* 1. Then 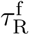 is equal to 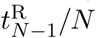 (see Eq. S30), with *γ* = 0. Using Eq. S30, we obtain

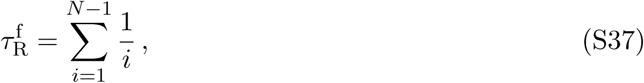

which entails 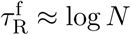 for N ≫ 1.

Finally, 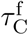 denotes the average time needed for the C mutants take over, starting from one C mutant and *N -* 1 R individuals. Considering R as type A and C as type B, we have *γ* = *f*_R_*/f*_C_ = *- δ*, independent whether antimicrobial is present or absent, and our initial condition is *j*_0_ = *N -* 1. Hence,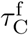 is given by 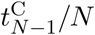 (see Eq. S30). In the neutral case *δ* = 0, it reduces to Eq. S22, and thus 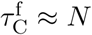 for *N ≫* 1. For *δ >* 0, it is shorter, as selection favors the fixation of C, and again a good approximation to the exact formula in Eq. S30 can be obtained within the diffusion approach [25] (see the Fokker-Planck equation below).

### 2. Large populations: deterministic limit

If stochastic effects are neglected, the dynamics of a microbial population can be described by coupled differential equations on the numbers of individuals of each genotype [25]. This deterministic approach is appropriate if the number *N* of competing microorganisms satisfies *Nµ*_1_ *≫* 1 [43]. Here, we derive and study the deterministic limit of the full-fledged stochastic model studied in the main text.

#### 2.1 From the stochastic model to the deterministic limit

Here, we present a full derivation of the deterministic limit of the stochastic model based on the Moran process (see above). This derivation closely follows those of Refs. [27, 28] and is presented here for the sake of pedagogy and completeness. Starting from the Master equation of our stochastic model, we obtain a Fokker-Planck equation, corresponding to the diffusion approximation [25], and then a deterministic differential equation, in the limits of increasingly large population sizes.

Let us first recall the Master equation corresponding to the Moran process, where *j* denotes the number of A individuals and *N - j* the number of B individuals, as above:

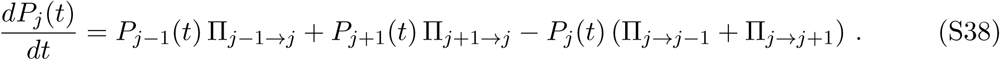

The notations in Eq. S38 are the same as in the previous section, and time is expressed in number of steps of the Moran process. Let us now introduce the reduced variables *x* = *j/N, τ* = *t/N*, as well as *ρ*(*x, τ*) = *NP*_*j*_(*t*). Then, since one step of the Moran process occurs each time unit, Eq. S38 can be rewritten as:

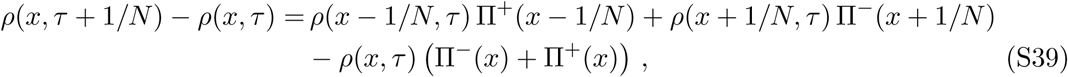

With

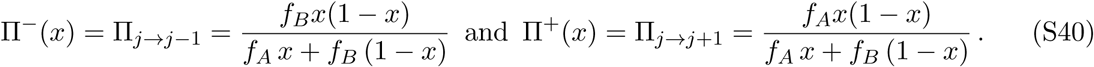

##### Diffusion approximation

For *N ≫* 1, considering that jumps are small at each step of the Moran process, i.e. 1*/N ≪x* and 1*/N ≪τ*, the probability density *ρ*(*x, τ*) and the transition probabilities ? ^*±*^ (*x*) can be expanded in a Taylor series around *x* and *τ*. This expansion, known as a Kramers-Moyal expansion [76], yields, to first order in 1*/N* :

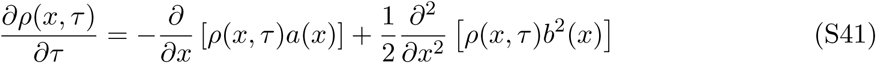

With

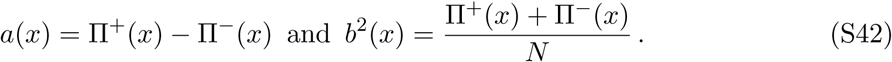

Eq. S41 is known as a diffusion equation, or a Fokker-Planck equation, or a Kolmogorov forward equation [76], and *a*(*x*) corresponds to the selection term (known as the drift term in physics), while *b*^2^(*x*) corresponds to the genetic drift term (known as the diffusion term in physics).

##### Deterministic limit

In the limit *N → ∞*, retaining only the zeroth-order terms in 1*/N*, Eq. S41 reduces to:

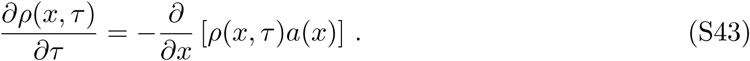

Let us focus on the average value of *x*, denoted by *(x)*. Using Eq. S41 yields

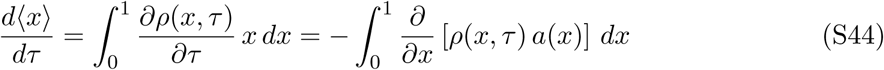

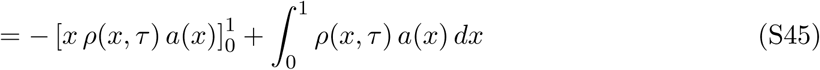

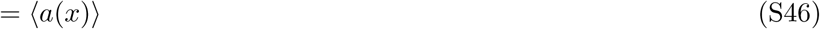

The first term of right hand side of Eq. S45 vanishes because *a*(0) = *a*(1) = 0. In the limit *N → ∞*, the distribution of *x* is very peaked around its mean, so ⟨*x⟩ ≈ x* and ⟨*a(x*) ⟩ ⟩ *≈ a*(*x*), yielding:

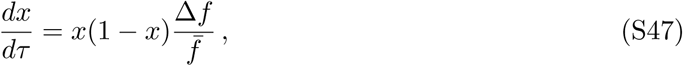

where Δ*f* = *f*_*A*_ *- f*_*B*_ denotes the difference of the fitnesses of the two types, while 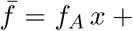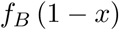 is the average fitness in the population. Eq. S47 is an ordinary differential equation known as the adjusted replicator equation [27]. Recall that *τ* corresponds to the number *t* of steps of the Moran process divided by the total number *N* of individuals in the population.

Hence, *τ* is the real time in numbers of generations used in the main text, and Eq. S47 is the proper deterministic limit for our stochastic process.

Note that in the framework of the Moran process, fitnesses are only relative. If one wanted to account for absolute fitness effects, so that a whole population reproduces faster if its average fitness is higher, one would need to include an additional rescaling of time 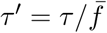, yielding a standard replicator equation:

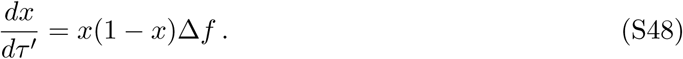

#### 2.2 Deterministic description of the evolution of antimicrobial resistance

##### System of ordinary differential equations

Let us now come back to our model of the evolution of antimicrobial resistance, with three types of microorganisms (see Fig. 1A). In the limit of large populations, the full-fledged stochastic model described in the main text will converge to a deterministic system of ordinary differential equations, as demonstrated above. Generalizing Eq. S48, by considering three types of individuals and taking into account mutations, yields a system of replicator-mutator equations [28]:

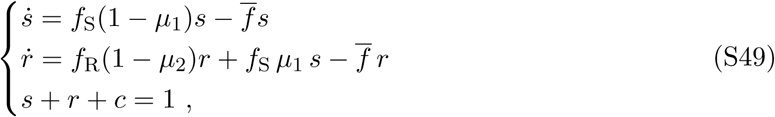

where *s, r* and *c* are the population fractions of S (sensitive), R (resistant) and C (resistant-compensated) microorganisms, respectively, while *f*_S_, *f*_R_ and *f*_C_ denote their fitnesses, *f* = *f*_S_ *s* + *f*_R_ *r* + *f*_C_ *c* denotes the average fitness in the population, and dots denote time derivatives. As demonstrated above, the deterministic limit of our stochastic model yields adjusted replicator equations (see Eq. S47). For the sake of simplicity, the present analytical discussion focuses on standard replicator equations (see Eq. S48). Recall that the correspondence can be obtained by a simple rescaling of time (see above).

The system of equations Eq. S49 only concerns population fractions, and constitutes the large-population limit *N → ∞* of our stochastic model at constant *N*. It is mathematically convenient to note that the same equations are obtained in the case of a population in which microorganisms have an exponential growth. This model, which enables to recover the system S49, is governed by the following system of linear differential equations:

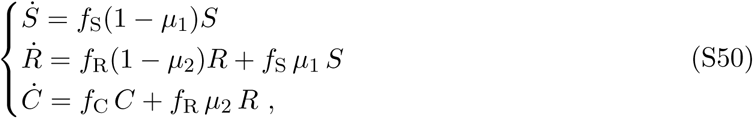

where *S, R* and *C* are the numbers of sensitive, resistant and resistant-compensated microorganisms, respectively.

##### Analytical resolution

Being linear, the system in Eq. S50 is straightforward to solve analytically:

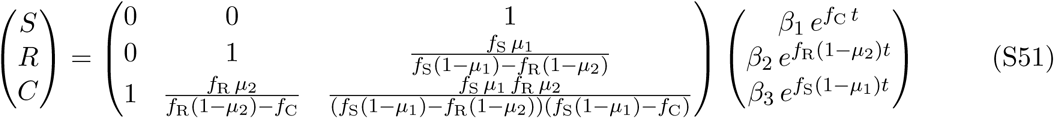

where *β*_1_, *β*_2_ and *β*_3_ can be expressed from the initial conditions *S*(0), *R*(0) and *C*(0). The fractions *s, r* and *c* can then be obtained from this solution, e.g. through *s* = *S/*(*S* + *R* + *C*).

##### Limiting regimes and characteristic timescales

As in the main text, we are going to focus on the case where the population initially only comprises sensitive microorganisms, i.e. *s*(0) = 1. In the case of periodic alternations of absence and presence of antimicrobial, a small fraction of R microorganisms will appear within the first half-period without antimicrobial. The subsequent evolution of the population composition can be separated into three successive regimes. In the first one, it suffices to consider S and R microorganisms, as the fraction of C is negligible, because the apparition of C requires an additional mutation. The second regime is more complex, and involves all three types of microorganisms, as the growth of C microorganisms makes the fractions of S and R microorganisms decrease. Then, provided that antimicrobial has been present for a sufficient time, the fraction of S microorganisms becomes negligible, because they cannot divide with antimicrobial. Hence, the third regime only involves R and C microorganisms, and does not depend on the presence or absence of antimicrobial, because the fitnesses of R and C are unaffected. Here, we determine analytically the main timescales involved in these first and third regimes.

##### First regime: S vs. R.

Let us consider the first regime where there are almost only S and R microorganisms. We are interested in the population fractions *s*(*t*) and *r*(*t*), with *s*(*t*)+*r*(*t*) *≈* 1. Eq. S49 then gives:

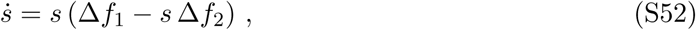

where we have defined Δ*f*_1_ = *f*_S_(1 *- µ*_1_) *- f*_R_(1 *- µ*_2_) and Δ*f*_2_ = *f*_S_ *- f*_R_(1 *- µ*_2_). Note that we expect Δ*f*_1_ *≈* Δ*f*_2_, since biologically relevant values generally satisfy *µ*_1_, *µ*_2_ ≪ 1 and *µ*_1_, *µ*_2_ ≪ *δ*. The solution of Eq. S52 reads

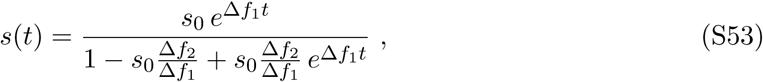

where *s*_0_ is the fraction of S microorganisms at the beginning of the first regime (taken as *t* = 0 here). In the presence of antimicrobial (*f*_S_ = 0), the previous expression can be simplified, using Δ*f*_1_ = Δ*f*_2_ = *-*(1 *- δ*)(1 *- µ*_2_). This allows us to identify the characteristic time *τ*_1_ of the decay of *s*, as R microorganisms take over:

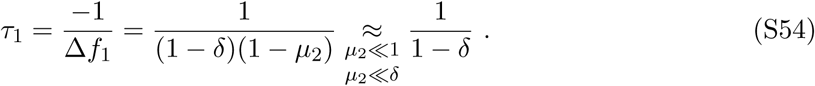

The duration *t*_1_ of the first regime in the presence of antimicrobial is governed by *τ*_1_. More precisely, Eq. S53 yields:

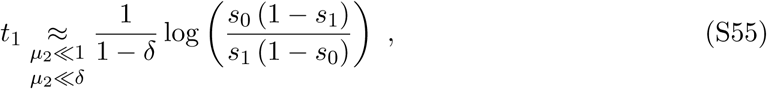

where *s*_1_ is the fraction of S microorganisms at the end of the first regime, at which point the fraction of C microorganisms is no longer negligible.

##### Third regime: R vs. C.

Let us now turn to the third regime, assuming that antimicrobial has been present for a long time enough to allow S microorganisms to become a small minority. Eq. S49 then gives:

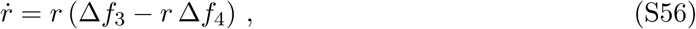

with Δ*f*_3_ = *f*_R_(1 *- µ*_2_) *- f*_C_ = *-δ*(1 *- µ*_2_) *- µ*_2_ and Δ*f*_4_ = *f*_R_ *- f*_C_ = *-δ*, independently of whether antimicrobial is present or not. Again, we generically expect Δ*f*_3_ *≈* Δ*f*_4_. The solution of Eq. S56 reads

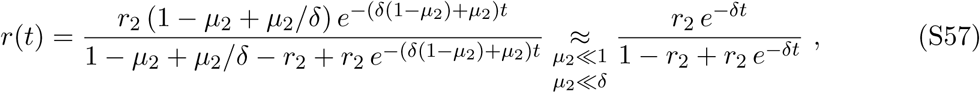

where *r*_2_ is the fraction of R microorganisms at the beginning of the third regime (taken as *t* = 0 here). Hence, the characteristic time *τ*_3_ of the decay of *r* reads:

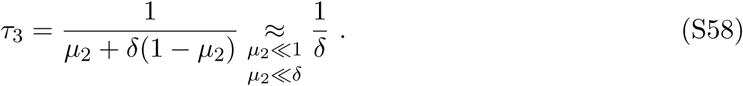

The duration *t*_3_ of the third regime in the presence of antimicrobial is governed by *τ*_3_. More precisely, Eq. S57 yields:

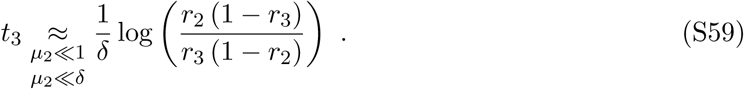

where *r*_3_ is the fraction of R microorganisms at the end of this regime, when C has become dominant in the population.

Note that the timescales obtained here are governed by selection (through the relevant fitness differences *δ* and 1 *- δ*). This stands in contrast with the results from our stochastic model (see main text) where mutation rates are crucial, especially through the waiting time before resistant mutants appear. In the deterministic description considered here, small fractions of resistant mutants appear right away, so this consideration is irrelevant. However, mutation rates come into play in the durations of the different regimes within the deterministic model, through the fractions of each type of microorganisms at the beginning and at the end of each regime, but with a weak logarithmic dependence (see Eqs. S55-S59).

#### 2.3 Comparison of stochastic and deterministic results

As in the main text, we now focus on the impact of a periodic presence of antimicrobial on the time it takes for a population to fully evolve resistance. For large microbial populations satisfying *N ≫* 1*/µ*_1_, we wish to check that the system of differential equations in Eq. S49 recovers the results obtained with our stochastic model. To this end, we solve the system in Eq. S49 numerically in the case of a periodic presence of antimicrobial. Note that complete fixation of a genotype does not happen in the deterministic model. Conversely, in the stochastic model, for a population of size *N*, the fixation of C corresponds to the discrete Moran step where the fraction *c* jumps from 1 *-* 1*/N* to 1. Hence, for our comparison between the deterministic results and the stochastic ones obtained for *N* microorganisms, we consider that C effectively fixes in the deterministic model when the fraction *c* reaches 1 *-* 1*/N*. In addition, for exactness, we use a numerical resolution of the system in Eq. S49 where time is rescaled through 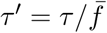. Indeed, the proper deterministic limit of our stochastic model corresponds to modified replicator equations, such as Eq. S47 (see above).

Fig. S2 shows that the deterministic model yields results very close to those obtained through the stochastic model, in the case of large population sizes *N ≥* 1*/µ*_1_. We recover the regimes described in the main text, with a plateau for short periods, and a linear dependence on *T* for larger ones. Moreover, the relative error made by using the deterministic model instead of the stochastic one is less than *∼* 20% (resp. *∼* 10%) for all data points with *N* = 10^5^ (resp. *N* = 10^6^) in Fig. S2.

Let us now present an analytical approximation for 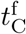, based on the different timescales computed previously. As the population is initially only composed of S microorganisms, they will remain dominant during the first half-period without antimicrobial, since they are fitter than R mutants (and we assume that *T/*2 is not large enough to extend to the point where C starts being important, which would then correspond to the valley crossing case). Afterwards, R microorganisms start growing fast during the second half-period. Note that in the deterministic case, there is always a nonzero fraction of resistant microorganisms at the end of the first half-period without antimicrobial, contrary to the stochastic case studied in the main text. Hence, we compute the fraction *s*_0_ = *s*(*T/*2) of S microorganisms at the end of the first half period, by using results for the above-described first regime without antimicrobials. This fraction *s*_0_ = *s*(*T/*2) is then taken as the initial condition of the first regime with antimicrobial. Then, for simplicity, we assume that *s* decays until it reaches *s*_1_ *≈* 0.1 (so *r*_1_ *≈* 0.9), while remaining in the first regime described above, in the presence of antimicrobial. We then assume the duration of the second regime is negligible, and consider that the third regime process starts right away, with a fraction *r*_2_ *≈* 0.9. As explained above, we consider that the third regime ends upon effective fixation of C, i.e. when *c* reaches 1 *-* 1*/N*, which implies *r*_3_ = 1*/N*. Using Eqs. S55 and S59, we obtain:

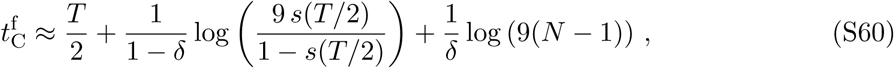

where *s*(*T/*2) is obtained by using Eq. S53 in the absence of antimicrobial:

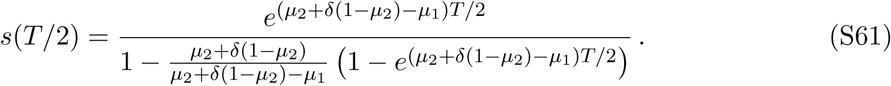

Eqs. S60-S61 yield good approximations of the analytical results obtained by numerical resolution of Eq. S49, as can be seen on Fig. S2. More precisely, the relative error made by using this approximation instead of the full numerical resolution is less than *∼* 13% for all parameters in Fig. S2.

For *T ≫* 2*/δ*, Eq. S61 reduces to *s*(*T/*2) *≈* 1 *- µ*_1_*/*[*µ*_2_ +*δ*(1 *- µ*_2_)] *≈* 1 *- µ*_1_*/δ*, so only the first term in Eq. S60 then depends on *T*. Hence, this term becomes dominant for large *T*, yielding 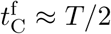 in this limit. This asymptotic behavior is again consistent with our predictions from the stochastic model (see main text). The horizontal purple solid line at large *T* in Fig. 3A, and the horizontal solid lines at large *N* in Fig. 3B, both correspond to 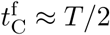, showing excellent agreement with our stochastic simulations as well.

Conversely, for small periods, the first term of Eq. S60 can be neglected, so the dependence on *T* of 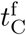 is weaker (Eq. S61 reduces to *s*(*T/*2) *≈* 1 *- µ*_1_*T/*2 for *T ≪*2*/δ*, so a weak logarithmic dependence on *T* remains, due to the second term of Eq. S60). It is interesting to note that the third term of 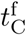 in Eq. S60 also increases logarithmically with *N*. This stands in contrast with the case of smaller populations, where our stochastic study showed that 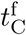 essentially decreases linearly with *N* (see main text). This change of behavior as *N* increases can be seen on Fig. 3A in the regime of small *T* (in particular, for large *N*, the purple data points corresponding to *N* = 10^6^ are then slightly higher than the blue ones corresponding to *N* = 10^5^; see also Fig. S2, where the y-axis range and scale are the same on panels A and B).

### 3. Robustness of the binary antimicrobial action model

Throughout our study, we have modeled the action of the antimicrobial in a binary way: below the MIC (“absence of antimicrobial”), growth is not affected, while above it (“presence of antimicrobial”), sensitive microorganisms cannot grow at all (see Model section in the main text). The relationship between antimicrobial concentration and microorganism fitness is termed the pharmacodynamics of the antimicrobial [62, 20]. Our binary approximation is motivated by the usual stiffness of pharmacodynamic curves around the MIC [20]. However, this stiffness is not infinite, and it is different for each antimicrobial. Here, we investigate the robustness of our binary model.

If one goes beyond the binary model and accounts for the smoothness of the pharmacodynamic curve, one additional factor enters the determination of the time dependence of fitness. It is the time dependence of the antimicrobial concentration, typically in a treated patient, which is known as pharmacokinetics [62, 20]. In fact, the time dependence of the fitness of sensitive microorganisms will be determined by a combination of pharmacodynamics and pharmacokinetics. Experimental pharmacodynamic curves are well-fitted by Hill functions, and pharmacokinetic curves are often modeled by exponential decays of drug concentration after intake [20]. The fitness versus time curve upon periodic antimicrobial intake will be a smooth periodic function resulting from the mathematical function composition of these two empirical relationships. The main feature of this curve will be how smooth or stiff it is, which can be characterized by its rise time, i.e. the time it takes to rise from *f*_S_ = 0.1 to *f*_S_ = 0.9 (if the fitness *f*_S_ of sensitive microorganisms ranges between 0 at very high antimicrobial concentrations and 1 without antimicrobial).

Thus motivated, we consider a smooth and periodic fitness versus time relationship *f*_S_(*t*) (see Fig. S3A), and we study the impact of the rise time Θ on the evolution of antimicrobial resistance in a microbial population. In practice, our smooth function, shown in Fig. S3A, is built using the error function erf 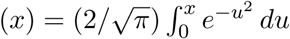, such that over each period of duration *T* :

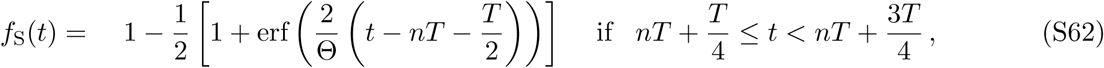

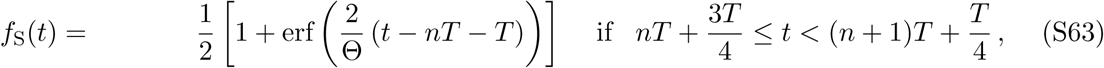

where *n* is a non-negative integer. In addition, we take *f*_S_(*t*) = 1 for 0 *≤ t ≤ T/*4, i.e. we start without antimicrobial at *t* = 0, and the first decrease of fitness occurs around *t* = *T/*2, in order to be as close as possible to our binary approximation (see Fig. 1B). Finally, as an extremely smooth case, we consider the case of a fitness *f*_S_ modeled by a sine function of period *T*, with the same initial condition and phase as our function with variable smoothness.

We have performed stochastic simulations using the model described in the main text, but with the fitness versus time relationship given in Eqs. S62-S63. Fig. S3 shows that for small rise times Θ, the dependence on the period *T* of the total time 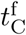 of full resistance evolution is the same as with our binary approximation, provided that the rise time is much smaller than the period, Θ ≪ *T*. Conversely, for small Θ satisfying Θ ≥ *T*, in which case our function is very smooth even though the absolute rise time is short, the behavior of 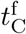 is similar to that obtained for the sine function. For larger values of Θ, namely Θ *≫* 10, the binary case is no longer matched when Θ ≪ *T*, and instead, a behavior intermediate between the binary case and the sine case is observed. This intermediate behavior gets closer to that observed in the sine case as Θ is increased.

These results can be rationalized as follows. When Θ is smaller than the relevant evolutionary timescales identified in the main text (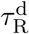, 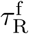 and 1/*μ*_1_), the shortest ones being 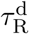 and 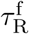 for *Nµ*_1_ ≪ 1), no relevant evolutionary process process can happen during a single smooth rise or decay of the fitness. If in addition Θ is much smaller than the environmental timescale *T*, then the fitness versus time function is stiff and effectively binary. However, if Θ is not much smaller than *T*, then the function is smooth, and the binary approximation is inappropriate. Finally, if Θ is longer than the shortest relevant evolutionary timescales (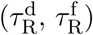), then relevant evolutionary processes can happen within a single smooth rise or decay of the fitness, and the behavior is more complex. In a nutshell, our binary approximation is appropriate provided that the rise time satisfies 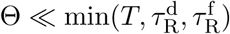.

